# An unbiased characterization of the HLA-E and CD94/NKG2x peptide repertoire reveals peptide ligands that skew NK cell activation

**DOI:** 10.1101/2022.08.03.502719

**Authors:** Brooke D. Huisman, Ning Guan, Timo Rückert, Lee Garner, Nishant K. Singh, Andrew J. McMichael, Geraldine M. Gillespie, Chiara Romagnani, Michael E. Birnbaum

## Abstract

HLA-E is a non-classical class I MHC protein involved in innate and adaptive immune recognition. While recent studies have shown HLA-E can present diverse peptides to NK cells and T cells, the HLA-E and NK receptor peptide repertoire has remained poorly defined, with only a limited number of peptide ligands identified. Here we screen a yeast-displayed peptide library in the context of HLA-E to identify 500 high-confidence unique peptides that bind both HLA-E and CD94/NKG2A or CD94/NKG2C. Utilizing the sequences identified via yeast display selections, we train prediction algorithms and identify human and cytomegalovirus (CMV) proteome-derived, HLA-E-presented peptides capable of binding and signaling through both CD94/NKG2A and CD94/NKG2C. In addition, we identify peptides which selectively activate NKG2C^+^ NK cells. Taken together, characterization of the HLA-E-binding peptide repertoire and identification of NK activity-modulating peptides present opportunities for studies of NK cell regulation in health and disease, in addition to vaccine and therapeutic design.

## Introduction

Human leukocyte antigen E (HLA-E) is a non-canonical, class Ib MHC expressed on most nucleated cells in the body. HLA-E has traditionally been thought to function by presenting sequences derived from a highly conserved segment from class I MHC (MHC-I) signal peptides for recognition by NK cells ^1,2^. Recognition of HLA-E in complex with these peptides serves as a mechanism to assess “missing self”: when pathogens downregulate class I MHC expression, HLA-E surface expression is in turn decreased, resulting in NK cell cytotoxic activity due to the loss of an inhibitory signal ^2,3^.

HLA-E’s cognate receptor heterodimers, the inhibitory CD94/NKG2A and the activating CD94/NKG2C (collectively referred to here as CD94/NKG2x), are expressed by NK cells and certain T cells. CD94/NKG2A is an ITIM motif-containing inhibitory receptor, while CD94/NKG2C is an ITAM motif-associating activating receptor ^4,5^. CD94/NKG2x bind HLA-E in a peptide-dependent manner ^6^, and NK cells integrate signaling from these and other receptors to modulate their cytotoxic activity ^7^. The fraction of NK cells expressing NKG2A and NKG2C varies widely across patients and disease states, with a large dependency on CMV serostatus ^8,9^. Moreover, NKG2C^+^ NK cells have also been associated with a lower viral setpoint in HIV infection ^9–11^ as well as with a reduced leukemia relapse in patients undergoing hematopoietic stem cell transplantation^12^, pointing towards strong anti-viral and anti-tumor activity of NKG2C^+^ NK cells. CD94/NKG2C and CD94/NKG2A are also expressed by significant fractions of tumor-infiltrating CD8^+^ T cells ^13,14^. Accordingly, modulation of NK and T cell function through CD94/NKG2x receptors is of therapeutic interest; indeed, NKG2A has been proposed as a possible therapeutically relevant immune checkpoint, and anti-NKG2A monoclonal antibodies alone or in combination promote anti-tumor immunity in mouse models and Phase II clinical trials ^15–17^.

Beyond the canonical MHC-I-derived peptide ligands, recent work suggests the HLA-E peptide repertoire is more diverse than previously thought, and has motivated study of TCR-mediated responses to HLA-E homologues in other species (MHC-E) ^18–20^. Vaccine strategies against SIV, HBV, and typhoid, including CMV-vector-based vaccines, elicit MHC-E-restricted CD8^+^T cell responses ^19,21–23^. Inducing HLA-E-restricted T cell responses may be a particularly attractive therapeutic strategy due to the essentially invariant nature of HLA-E as compared to class Ia MHCs, allowing for a given peptide therapy to be potentially beneficial for all individuals.

Despite interest in HLA-E and CD94/NKG2x for infectious disease and cancer treatment, a limited number of HLA-E peptide binders are currently known, with approximately 700 total contained in the Immune Epitope Database (IEDB) ^24^, but likely do not represent a systematic assessment of all possible HLA-E binders. Current state-of-art MHC motif predictors such as NetMHC ^25^ and MHCflurry ^26^ may therefore be limited in HLA-E prediction performance due to the scarcity of available training data.

Here, we used a yeast display-based approach to conduct high-throughput analysis of the HLA-E and CD94/NKG2x peptide repertoire. Starting with a randomized library of 100 million unique peptides linked to HLA-E, we selected for binding to HLA-E and either CD94/NKG2A or CD94/NKG2C. Deep sequencing of the HLA-E selection libraries identified ~500 high-confidence unique peptide binders. With these data, we developed computational algorithms to predict proteome-derived peptide binders to HLA-E and NK receptors. To validate a set of potential peptide binders, we performed biophysical validation, including peptide stability assays, surface plasmon resonance (SPR), and NK cell activation analysis. We identify human- and CMV-derived peptides able to affect NK cell effector functions, including peptides which selectively lead to NK cell activation. This improved understanding of the HLA-E and NK receptor repertoires could serve as a tool for therapeutic design, and has demonstrated utility for identification of peptides capable of skewing NK cell activity.

## Results

### Design of HLA-E for yeast display and library selections using CD94/NKG2x

To study the diversity of the HLA-E peptide repertoire, we adapted an MHC yeast surface display system previously used to study class Ia MHC-restricted TCR recognition ^27,28^. In this system, peptide, Beta-2-microglobulin (β2M), and MHC heavy chain are covalently linked to each other and fused to the N-terminus of the yeast mating factor protein Aga2p (**Figure 1A, 1B**). The HLA-E single chain trimer construct expressed well on the yeast surface, as confirmed via staining for an included epitope tag and via an anti-HLA-E antibody (**Figure 1C**). Since epitope tag staining assesses construct expression rather than protein fold ^29–31^, we tested peptide-HLA-E (pHLA-E) fold by staining with its cognate NK receptors. Using a canonical signal peptide-derived sequence (VMAPRTLFL, “VL9”), pHLA-E stained with CD94/NKG2A tetramer showed a positive population, indicating pHLA-E was correctly folded for receptor recognition (**Figure 1C, Supplemental Figure 1A**). CD94/NKG2C tetramer also stained VL9-expessing yeast but with lower signal, consistent with its lower affinity (**Supplemental Figure 1B**).

**Figure 1.**
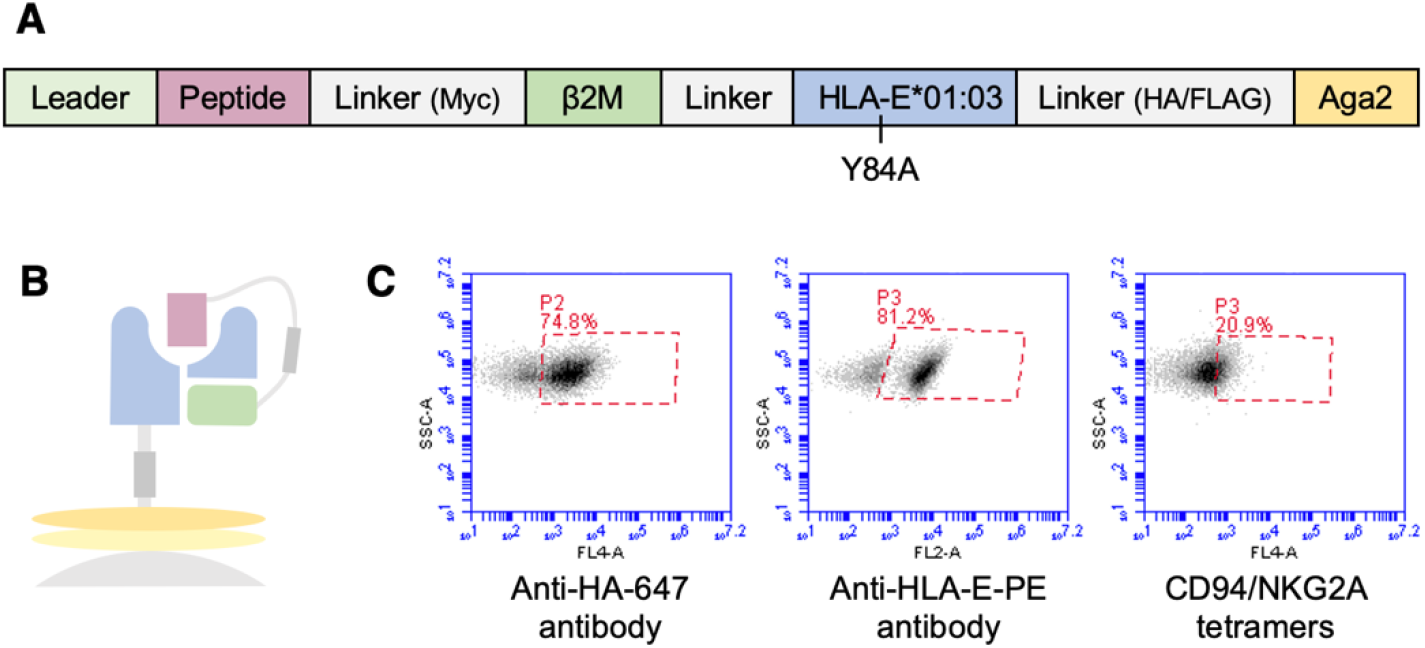
Design and validation of HLA-E on yeast. Representation of HLA-E **a)** sequence and **b)** on the surface of yeast. **c)** Validation staining of HLA-E with VMAPRTLFL peptide by anti-HA-647 and anti-HLA-E-PE antibodi**es** and with CD94/NKG2A tetramers made with Streptavidin-647.

Next, we created a 9mer peptide library using the pHLA-E construct. To screen for peptide preferences for both HLA-E and CD94/NKG2x binding, we fully randomized each position along the peptide rather than limiting sequence diversity at the P2/P9 anchor positions as described for previous pMHC libraries^27^. We then iteratively enriched the randomized peptide library displayed by HLA-E for binding to CD94/NKG2A or CD94/NKG2C via magnetic selection (**Supplemental Figure 2A**). After three rounds of selecting with biotinylated recepto**rs** bound to streptavidin-coated beads, we conducted both iterative and cross-selections using CD94/NKG2x tetramers, which were more stringent due to tetramers’ lower avidity (**Supplemental Figure 2B**). Cross-selection was performed to study ability of NKG2C-enriched peptides to bind NKG2A, and vice versa. CD94/NKG2A showed similar tetramer staining for yeast previously selected by CD94/NKG2A or CD94/NKG2C for three rounds, indicating that the peptide binding motifs of CD94/NKG2A and CD94/NKG2C may have significant similarities (**Supplemental Figure 2B**), consistent with sequence conservation between NKG2A and NKG2C at HLA-E- and peptide-contacting residues ^32^.

### Analysis of library-enriched peptides reveals highly diverse peptides with much overlap between CD94/NKG2A and CD94/NKG2C repertoires

Following selections, we performed deep sequencing of each round of library-enriched yeast to determine the identities of the enriched peptides (**Supplemental Data 1**). We identify HLA-E 9mer library hits with significantly greater diversity than the canonical MHC-I leader-like ligands ^18,20^. Specifically, deep sequencing showed ~10^4^ unique peptides per sample after the third round, and ~5 x10^3^ unique peptides per sample after the fourth round of selection, although approximately 500 peptides are dominant among the enriched peptides, each with read counts >100 (**Supplemental Figure 3**).

A clear motif of enriched peptides emerged in the data after the third round of selection (**Figure 2**, **Supplemental Figure 4**). The motif, highly conserved between CD94/NKG2A-selected and CD94/NKG2C-selected libraries, contained significant diversity on the peptide N-terminus. Among primary anchors ^33^, P2 accommodated multiple hydrophobic residues, and P9 strongly preferred Leu. P3 enriched strongly for Pro, with a stronger preference than the traditional P2 anchor residue preferences. As secondary anchors, P6 enriched for Ser and Thr, and P7 enriched for Leu in addition to other hydrophobic residues. We generally observed greater conservation of the peptide C-terminus (P5 through P9) in enriched sequences, likely corresponding to constraints imposed by the CD94/NKG2x-pHLA-E binding interface^32,34^. P5 and P8 are primary CD94/NKG2x-contact residues ^35^. Our selection data identified P5 with strong preference for Arg, while P8 displayed preference for Trp, Phe, or Leu. CD94/NKG2A demonstrated more relaxed binding constraints as compared to CD94/NKG2C, potentially due to CD94/NKG2A’s higher affinity for HLA-E ^32^. While the motifs after Round 4 selection with CD94/NKG2x tetramers were consistent with Round 3 motifs, CD94/NKG2A tetramer selection led to a less diverse motif likely containing higher affinity binders (**Supplemental Figure 4**). Through peptide clustering, we also observed a subset of peptides characterized by strong preference for P1 Trp and P2 Asn (“WN peptides”) with amino acid preferences at the N-terminus of the peptide yet little constraint on the C-terminal portion of the peptide (**Supplemental Figure 5**).

**Figure 2.**
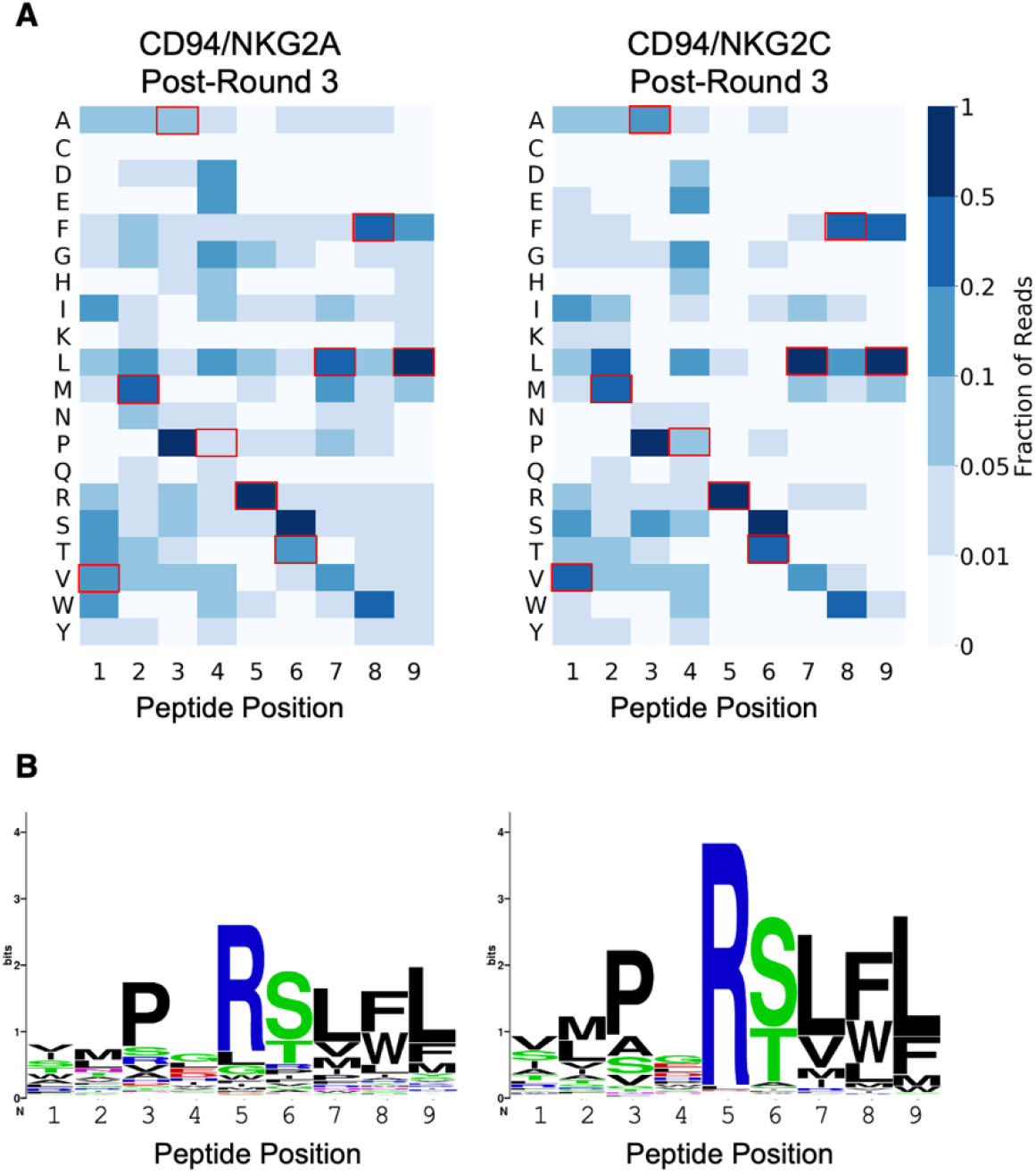
Peptide repertoire of HLA-E with CD94/NKG2A or CD94/NKG2C. **a)** Heatmaps showing peptide positional amino acid preferences for binding HLA-E and CD94/NKG2A or CD94/NKG2C with peptides weighted by read count. Residues corresponding to VL9 peptide are highlighted with red boxes. **b)** Sequence logos present the same data, also weighted by read count, generated with WebLogo.

While the majority of described HLA-E-restricted peptides have thus far been 9 amino acids long ^24^, class Ia MHCs can bind a range of additional peptide lengths, with 10mers being nearly as prevalent as 9mers for some alleles ^36^. Therefore, in addition to the 9mer library, we also generated at 10mer peptide library linked to HLA-E and performed selections with CD94/NKG2C. Interestingly, the motif for the first 9 amino acids of the 10mer largely resembled the motif in the 9mer library (**Supplemental Figure 6**). The C-terminal amino acid showed less preference than the 9mer library PΩ position, consistent with the 10mer peptides binding as 9mer peptides, with the tenth residue acting as part of the peptide linker for the single chain trimer-formatted pMHC. This suggests that 9mer peptides preferentially bind to CD94/NKG2x in the context of HLA-E compared to 10mer peptides, leading us to focus on the 9mer library data.

### Training prediction algorithms and predicting human- and pathogen-derived ligands

Given the large theoretical space (20^9^) of 9mer peptides, yeast displayed peptide libraries do not comprehensively examine all possible peptide sequences. We therefore trained a prediction algorithm using our yeast display-derived peptide data to predict putative human- and pathogen-derived peptide binders to HLA-E and CD94/NKG2x. We used yeast selection data to train the NNAlign architecture ^37^, which we have previously shown produces high-confidence class II MHC-peptide binding predictions using yeast display-derived data ^31^. We trained separate models using yeast display data from Round 3 of CD94/NKG2A and CD94/NKG2C selections, with negative examples drawn from the unselected library.

We applied this algorithm to the 11 million unique 9mers, generated from a sliding window of size 9, step size 1, along a reference human proteome. Given the absence of relevant comparator methods for predicting HLA-E and CD94/NKG2x binding, we examined the rankings of known binders. The ranks of the following canonical signal peptide-derived HLA-E ligands were examined: VMAPRTLFL, VMAPRTLLL, VMAPRTLIL, VMAPRALLL, VMAPRTVLL, VMAPRTLVL ^38–45^, with their predicted ranks shown in **Table 1**. The distribution of ranks for these peptides were significantly better than the human proteome as a whole (with *p*-value of 1.1 x 10^-5^ by Mann-Whitney U test, for either CD94/NKG2A and CD94/NKG2C model predictions).

**Table 1.** VL9 and VL9-like peptide predicted ranks. Predicted ranks of eight signal peptides by CD94/NKG2A and CD94/NKG2C models, ranked out of 11 million human-derived 9mer peptides.

| Peptides | NKG2A Model:<br>Rank | NKG2C Model:<br>Rank |
| --- | --- | --- |
| VMAPRTLFL | 224 | 2 |
| VMAPRTLLL | 418 | 12 |
| VMAPRTVLL | 547 | 22 |
| VMAPRTLIL | 481 | 32 |
| VMAPRTLVL | 602 | 48 |
| VMAPRALLL | 514 | 69 |

The top ten predicted human proteome-derived peptides are shown in **Table 2**. These binders largely share features of known binders, including hydrophobic P7-P9 residues and P5 Arg. One of the top predicted binders resembles the N-terminal motif of the subdominant WN peptides (UBAC2_275-279_). Proteins containing peptides such as BFAR_263-271_ (VNPGRSLFL; derived from bifunctional apoptosis regulator) and CREB3L1_419-427_ (QMPSRSLLF; derived from cyclic AMP-responsive element-binding protein 3-like protein 1) are associated with regulation of cell survival ^46–48^, and CREB3L1-expression has been reported to promote metastasis in certain cancers ^49^. MHC-E has been associated with stress response, and these peptides may represent potential sets of peptides HLA-E can present in states of cellular distress ^50^.

**Table 2.**
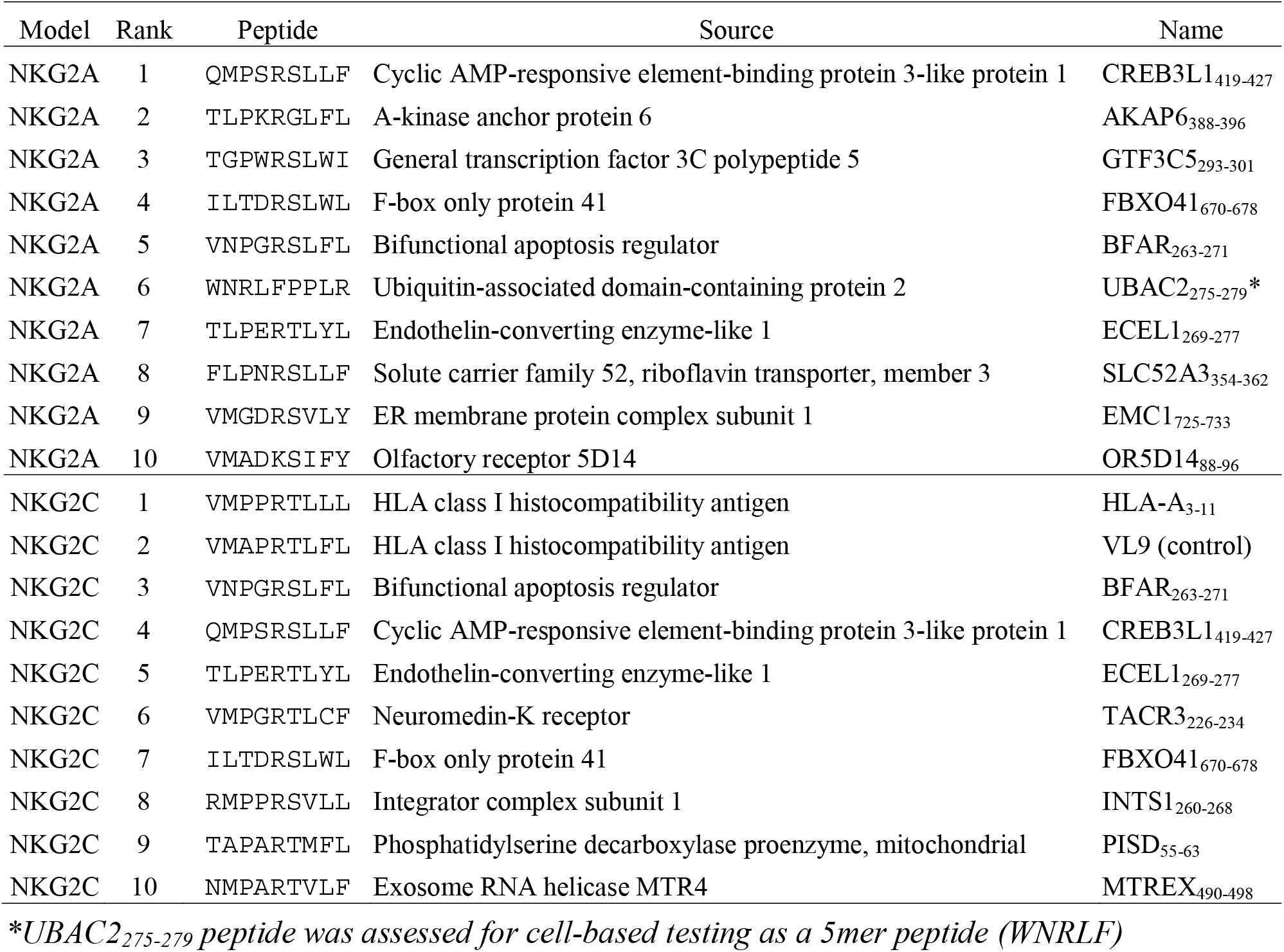
Top ten predicted human peptides from both models. Top predicted peptides by CD94/NKG2A and CD94/NKG2C models, with source proteins noted. *UBAC2_275-279_ peptide was assessed for cell-based testing as a 5mer peptide (WNRLF)

In addition to predicting on human-derived peptides, we applied our models to a reference human CMV proteome. One mechanism by which CMV evades T cell and NK cell detection is by generating a UL40-derived peptide identical to canonical VL9 and VL9-like signal peptides, which can be loaded onto HLA-E, even when canonical class I processing is disrupted ^38,45^. We ranked this UL40-derived peptide highly among CMV-derived peptides (**Table 3**). Interestingly, however, both CD94/NKG2A and CD94/NKG2C models rank a UL120-derived peptide VLPHRTQFL (UL120_72-80, Merlin_) more highly, which bears the hallmarks of a strong binder, including P3 Pro, P5 Arg, P8 Phe, and P9 Leu. Our predicted peptide UL120_72-80_, Merlin is from a highly polymorphic region among different CMV strains. For example, despite a large degree of conservation in the UL120 proteins between strain Merlin (used in our search) and strain AD169, strain AD169 has the divergent peptide SAPLKTRFL (“UL120_71-79, AD169_”) at the corresponding position.

**Table 3.** Top predicted CMV peptides from both models. Top predicted peptides by CD94/NKG2A and CD94/NKG2C models, with source protein noted. Ranks are among 61 thousand CMV-derived peptides.

| NKG2A<br>Model: Rank | NKG2C<br>Model: Rank | Peptide | Source | Name |
| --- | --- | --- | --- | --- |
| 1 | 1 | VLPHRTQFL | Membrane protein UL120 | UL120<sub>72-80</sub>, Merlin |
| 2 | 3 | TGAARSFFF | DNA helicase/primase complex-associated protein | UL102<sub>292-300</sub> |
| 3 | 2 | VMAPRTLIL | UL40 | UL40<sub>15-23</sub> |

### Validating peptides for HLA-E binding ability

To probe top-predicted proteome-derived peptides (**Table 2** and **Table 3**) for HLA-E binding, we assessed their ability to stabilize surface-expressed HLA-E on RMA-S/HLA-E cells, a TAP-deficient mouse tumor cell line engineered to express HLA-E and human β2M (**Figure 3, Supplemental Figure 7**; peptides with similar stabilization effects are plotted in the same color) ^51^. Ten human peptides showed strong stabilization of HLA-E (Human peptides INTS1_260-268_, HLA-A_3-11_, ECEL1_269-277_, TACR3_226-234_, CREB3L1_419-427_, AKAP6_388-396_, MTREX_490-498_, FBXO41_670-678_, SLC52A3_354-362_, PISD_55-63_). An additional two human peptides demonstrated weak but detectable stabilization (Human peptides BFAR_263-271_, GTF3C5_293-301_). Of the CMV peptides, UL120_72-80_, _Merlin_ showed strong HLA-E binding, and peptide variants in different viral strains (SAPLKTRFL “UL120_71-79, AD169_”; SVPLKTRFL “UL120_71-79, BE/33/2010_”) showed weaker but detectable stabilization. Results are consistent with thermal stability measurements taken on a subset of human and CMV proteome-derived predicted peptides (**Supplemental Figure 8**), with relative melting temperatures between peptides matching relative cell surface stabilization effects. Several peptides showed no stabilizing effects (Human peptides UBAC2_275-279_, EMC1_725-733_, OR5D14_88-96_, and CMV peptide UL102_292-300_), potentially due to disfavored residues that are unique to non-binders, such as P7 Phe and P9 Tyr (**Supplemental Figure 9**). We additionally tested peptide variants containing the enriched N-terminal ‘WN’ motif, which showed no detectable stabilization (**Supplemental Figure 9**).

**Figure 3.**
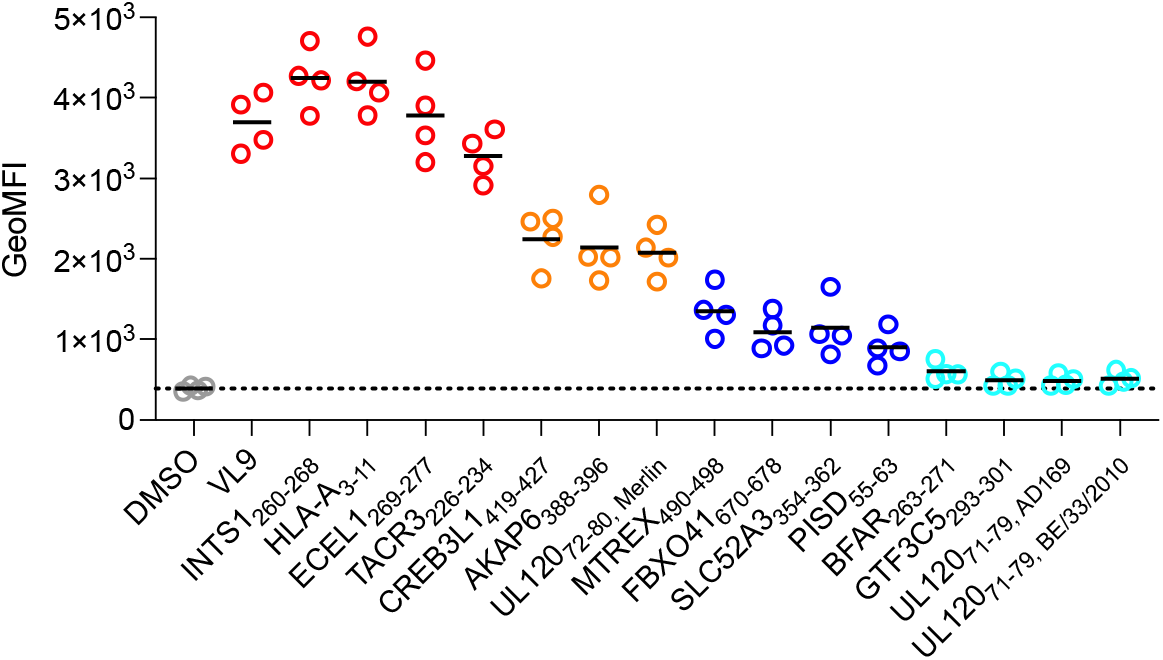
Peptide stabilization of HLA-E surface expression. Assessment of peptide-HLA-E binding via HLA-E surface stabilization assay with RMA-S/HLA-E cells incubated with 30 μM peptide. HLA-E expression is detected with an anti-HLA-E antibody. Measurements from replicate experiments are shown, with solid black lines indicating mean values.

### Proteome-derived peptides demonstrate ability to modulate NK cell activation

We hypothesized that our predicted proteome-derived peptides could affect NK cell activation through interactions with CD94/NKG2A and CD94/NKG2C. To investigate, we performed cell-based activation assays on primary NK cells (**Figure 4A**). NK cells were co-incubated with peptide-pulsed HLA-E-expressing K562 cells, as described previously ^51^. The parental K562 human myelogenous leukemia cell line is a typical target to measure NK cell cytotoxicity and activation ^51^, allowing for assessment of both NKG2A-mediated inhibition and NKG2C-mediated activation. All peptides which bound and stabilized HLA-E in the cell-based stability experiment (**Figure 3**) were analyzed for their effects on NKG2A^+^/NKG2C^-^ or NKG2A^-^ /NKG2C^+^ NK cell activity, as measured by their ability to induce NK cell degranulation as well as IFN-γ, TNF and CCL3 production, in comparison to class I MHC-derived positive control peptides VMAPRTLIL (LIL) and VMAPRTLFL (LFL), and negative control peptide VMAPQSLLL (PQS) (**Figure 4**; with coloring as in **Figure 3**) ^51^. NKG2A^-^/NKG2C^-^ NK cells served as internal control as these should be unaffected by peptide-HLA-E complexes (**Supplemental Figure 10**). Six peptides had both strong inhibitory effects on NKG2A^+^ cells and strong activating effects on NKG2C^+^ cells (Human peptides INTS1_260-268_, HLA-A_3-11_, ECEL1_269-277_, TACR3_226-234_, MTREX_490-498_; CMV peptide UL120_72-80, Merlin_), and four peptides had more moderate inhibitory effects on NKG2A^+^ cells while maintaining strong activating effects on NKG2C^+^ cells (Human peptides CREB3L1_419-427_, AKAP6_388-396_, FBXO41_670-678_, SLC52A_3354-362_). Strikingly, three peptides showed minimal inhibitory effects but clear activating effects (Human peptides PISD_55-63_, BFAR_263-271_, GTF3C5_293-301_). The UL120_71-79, AD169_ peptide seemed to negatively affect activation of all subsets, including the internal NGK2A^-^NKG2C^-^ control, suggesting nonspecific effects. UL120_71-79, BE/33/2010_ peptide showed minimal effects on both NKG2A^+^ or NKG2C^+^ cells.

In contrast to K562 cell lines which are targets for human NK cells, RMA-S cells (a TAP-deficient mouse tumor cell line) ^52^ are inert to human NK cells, unless engineered to express an immune molecule such as HLA-E and pulsed with appropriate peptides. Because of this, RMA-S/HLA-E cells have been shown to provide additional granularity on the strength of NKG2C-mediated NK cell activation ^51^. We performed additional NK cell activation assays with RMA-S/HLA-E cells (**Figure 4B**). Minimal activation is observed for the negative control peptide (PQS) and differential activation is observed for positive control LIL and LFL peptides, consistent with previous data ^51^. In previous studies, LIL and LFL peptides have demonstrated differential activating effects, with LFL peptide acting as the most potent activating peptide, which we observe as well (**Figure 4B**) ^51^. Remarkably, two peptides (Human peptides AKAP6_388-396_, FBXO41_670-678_) showed activation at the level of the strongest positive control binder (LFL) (**Figure 4B**). Additional peptides showed activation similar to the weaker positive control (LIL) or between the two positive controls (Human peptides HLA-A_3-11_, ECEL1_269-277_, TACR3_226-234_, CREB3L1_419-427_, SLC52A3_354-362_, BFAR_263-271_; CMV peptide UL120_72-80, Merlin_). Three peptides showed detectable activation, though below both positive control examples (Human peptides MTREX_490-498_, PISD_55-63_, GTF3C5_293-301_). In sum, among our predicted peptides, we have identified sequences able to modulate NK cell activity at the level of positive control ligands, and have additionally identified peptides that are able to selectively activate NK cells, with minimal inhibitory effects.

### Measuring pHLA-E-CD94/NKG2x affinity for identified peptides

To investigate the mechanism by which our activity-skewed peptides selectively activate via CD94/NKG2C without inhibiting through CD94/NKG2A, we measured the affinity of peptide-HLA-E for NK receptors via SPR (**Figure 5**). Specifically, we measured CD94/NKG2A and CD94/NKG2C affinity for HLA-E complexed with three sets of peptides. The first set are the most activity-skewed peptides (Human peptides GTF3C5_293-301_, BFAR_263-271_, and PISD_55-63_) which exhibited NKG2C-mediated activation and minimal NKG2A-mediated inhibition (**Figure 4**). The second set of peptides (Human peptides CREB3L1_419-427_ and AKAP6_388-396_) exhibited both activation and inhibition, through more modest inhibition compared to positive control peptides (**Figure 4**), as well as greater HLA-E stability than Human peptides GTF3C5293-301, BFAR_263-271_, and PISD_55-63_. Third, we included VL9 peptide as a positive control. All six peptides assessed exhibited higher affinity for CD94/NKG2A than CD94/NKG2C, suggesting that selective activation was not due to higher affinity for CD94/NKG2C (**Figure 5A** and **Figure 5C**). Further, the selectively activating peptides BFAR_263-271_ and GTF3C5_293-301_ were the highest and lowest affinity peptides for both receptors, respectively, suggesting that selective activation was also not due to an affinity threshold for signaling that differed between the two receptors. However, when comparing maximal response units (RU) to the theoretical Rmax, (**Figure 5B**), we observe GTF3C5_293-301_ and BFAR_263-271_ reach lower maximal values, including at saturation (BFAR_263-271_), suggesting that these peptide-HLA-E complexes are less stable, consistent with their HLA-E stabilization ability (**Figure 3**).

**Figure 4.**
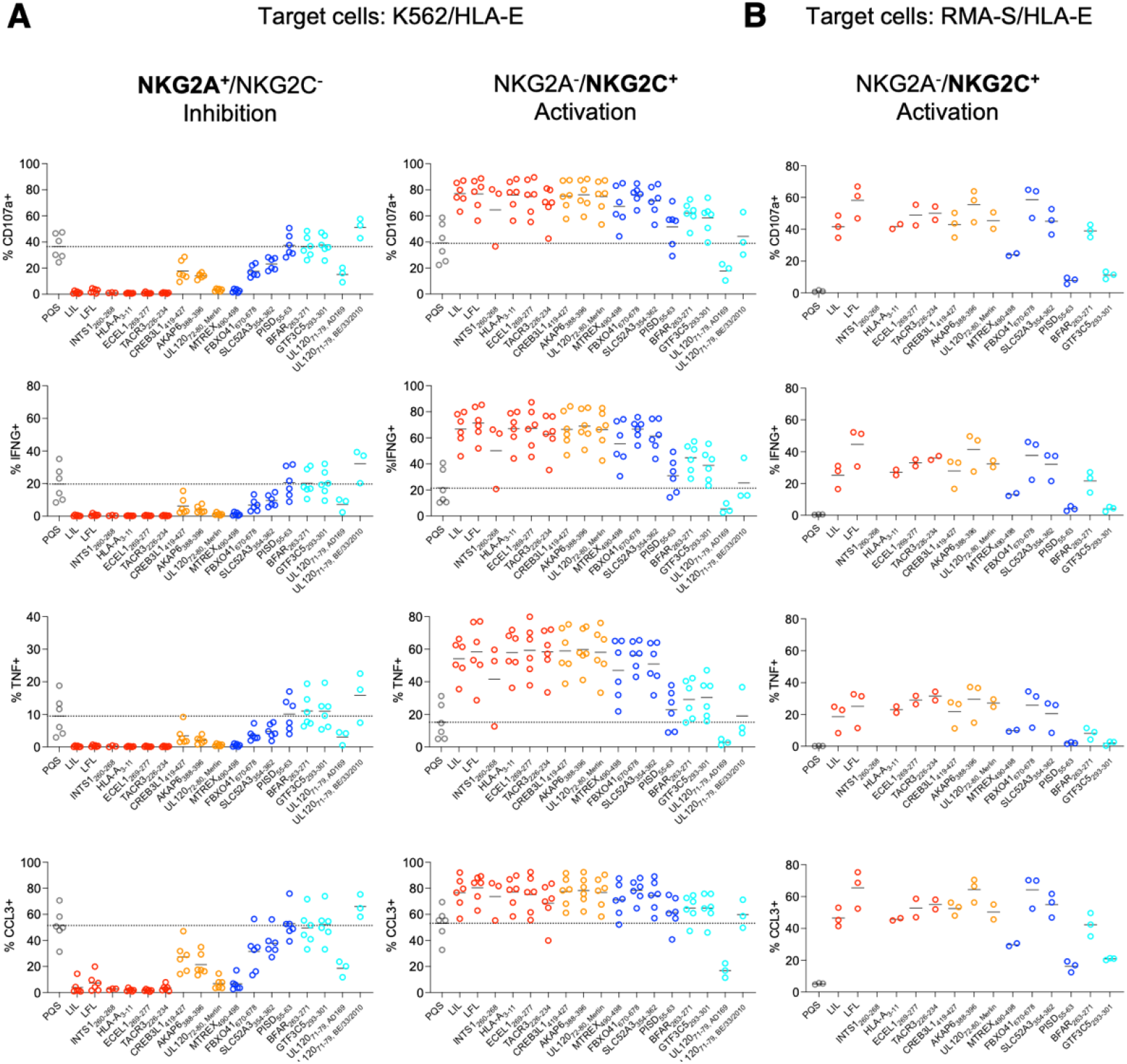
NK cell stimulation assays. Assessment of effects of peptides on NK cell activity through incubation of NK cells, peptides, and **a)** K562/HLA-E or **b)** RMA-S/HLA-E target cells. Peptides were derived from the human or CMV proteomes or are included as positive (LFL, LIL) or negative controls (PQS). Replicates for individual peptides are from different donors, with solid black lines indicating mean values.

**Figure 5.**
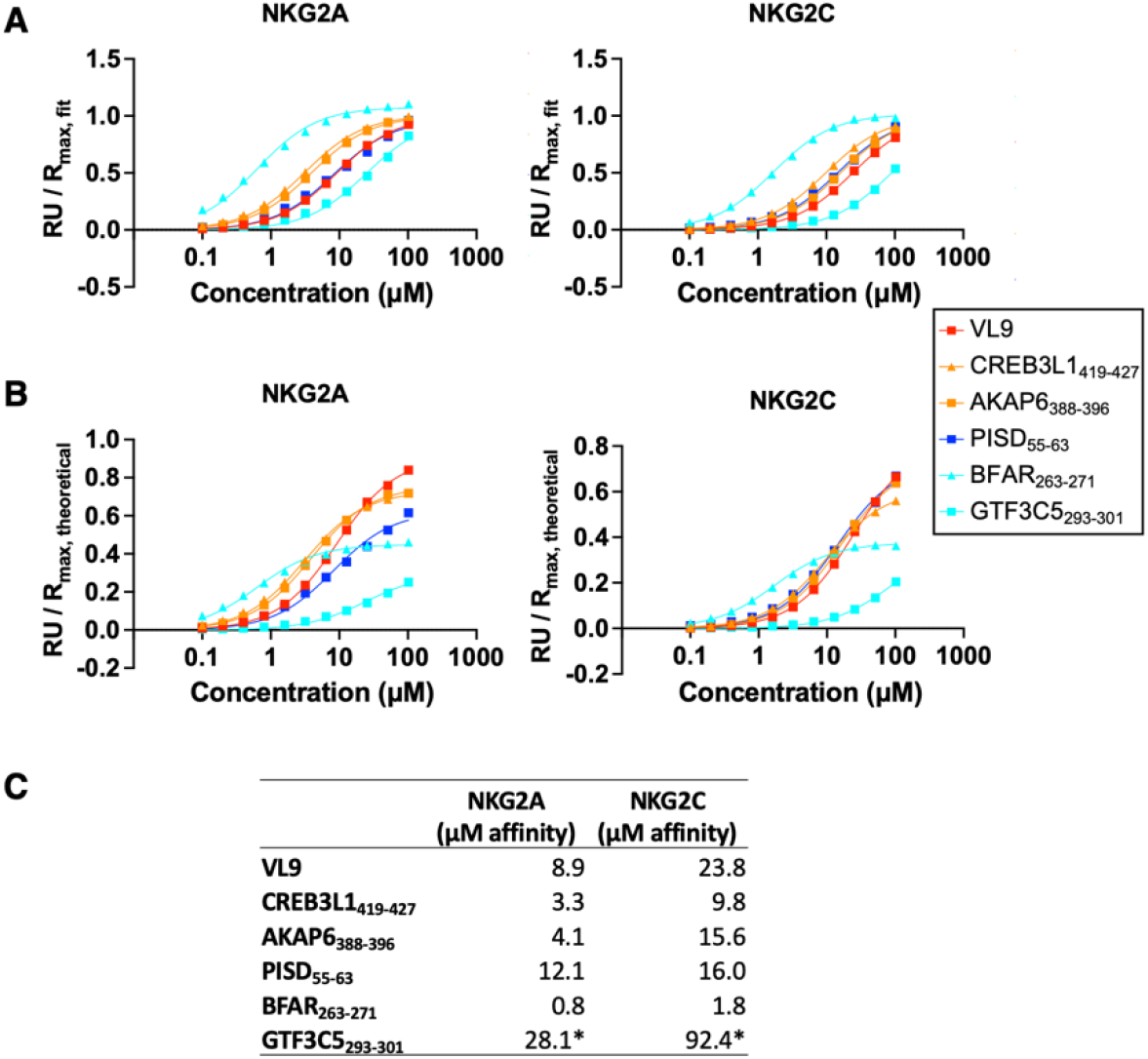
SPR of proteome-derived peptides with CD94/NKG2A and CD94/NKG2C. Affinity measurements of peptide-HLA-E for CD94/NKG2A and CD94/NKG2C, **a)** normalized to fitted R_max_ or **b)** theoretical R_max_. **c)** Summary of approximate K_D_ affinity values for CD94/NKG2x binding to peptide-HLA-E. *Indicates weak binders not approaching saturation, so K_D_ values are inexact.

We additionally assessed peptides containing the N-terminal WN motif for their ability to bind CD94/NKG2x in the context of HLA-E. Though WN peptides showed minimal ability to stabilize cell-expressed HLA-E (**Supplemental Figure 9**), a WN peptide expressed in a single chain trimer format with HLA-E was able to bind to CD94/NKG2A and CD94/NKG2C (**Supplemental Figure 5**), suggesting that the observed enrichment in the yeast display results was a bona fide interaction, but that the peptide-MHC interaction was sufficiently weak as to require covalent linkage to detect CD94/NKG2x binding.

## Discussion

HLA-E has been shown to present diverse peptides to NK and T cell receptors ^19,21,23^. This potential for peptide diversity, coupled with its sequence conservation across global populations^53^, has led to therapeutic interest in HLA-E and its cognate receptors ^13,15,17,21,23^. Despite this interest, the HLA-E peptide repertoire has remained poorly characterized. Here, we utilized yeast display libraries to characterize the repertoire of ligands that can bind to CD94/NKG2x in the context of HLA-E. Utilizing these data, we trained prediction algorithms and identified human- and CMV-derived peptides which can alter NK cell activity through CD94/NKG2x when presented by HLA-E. While many peptides can both activate through CD94/NKG2C and inhibit through CD94/NKG2A, a subset of peptides selectively activate CD94/NKG2C^+^ NK cells. Interestingly, these selectively activating peptides exhibited less HLA-E stabilization while showing comparable affinity to CD94/NKG2x, suggesting that altering peptide affinity to HLA-E, rather than the pHLA-E affinity to CD94/NKG2x, may be a mechanism for modulating NK cell activity. These effects may be due to potential differences in the level of receptor occupancy required to induce CD94/NKG2A vs CD94/NKG2C signaling, which may be linked to differences in the receptors’ signaling apparatuses ^5^.

Past characterization of the HLA-E peptide repertoire has varied widely, including primary anchor residue locations and preferences ^54^ and secondary anchor residue preferences ^35^, likely limited in part by study throughput. Our dataset of HLA-E and CD94/NKG2x-binding peptides is consistent with known high-confidence HLA-E binders and structural knowledge of the interface ^32^, while further characterizing the peptide repertoire. Notably, we identify a strong preference for Pro at position 3, which VL9 contains at position 4 and has been reported to be preferred at positions 3, 4, 6, or 7 ^33^. Other recent work includes a study from Ruibal et al, utilizing a combinatorial peptide library for screening peptides for HLA-E binding ^55^ which allows for the interrogation of peptide preferences, although it does not determine the identity of individual peptides. The yeast display approach allows us to determine the identities of individual peptides, which were useful here as training examples for prediction algorithms. Our datasets match at several positions, such as P9, where we both identify Phe and Leu as the two most preferred residues. However, important differences between the characterizations of HLA-E-binding residues are also present. For example, Ruibal et al identified a strong preference for P2 Met. In contrast, our dataset suggests a less extreme preference, supported by HLA-E binding of peptides which include P2 Leu, Asn, and Ala, and a more dominant preference for P3 Pro.

Using our yeast display-generated data, we trained algorithms to predict proteome-derived peptides for binding HLA-E and CD94/NKG2x. While there exist no algorithms for predicting peptide binding to both HLA-E and CD94/NKG2x, NetMHCpan4.0 can be utilized to predict peptide-HLA-E binding. NetMHCpan4.0 predicts that all of our proteome-derived 9mer peptides assessed in **Figure 3** and **Supplemental Figure 9** will bind to HLA-E, either as strong or weak binders ^25^, including peptides which showed no HLA-E stabilizing effects.

Newly characterized proteome-derived peptides which affect NK cell activity may present therapeutic opportunities. The identified CMV-derived peptide from the UL120 protein can alter NK cell activity, expanding the known CMV peptides able to bind HLA-E and CD94/NKG2x beyond the known UL40-derived peptide ^45^. Additionally, because related CMV strains are highly dissimilar at this region, with these peptides demonstrating different functional effects, we hypothesize that these differences between strains could lead to differential HLA-E presentation of CMV-derived peptides and may contribute to varying responses to CMV strains ^51,56^. Future work may investigate the relationship between UL120 protein sequence, NK cell responses, including adaptive NKG2C^+^ NK, and CMV viral titers, as well as investigate the existence of T cells reactive to these peptides in the context of HLA-E. Similarly, our studies on the human proteome reveal an unappreciated complexity of self-peptides capable of stabilizing HLA-E and modulating NK cell responses. Identified peptides are derived from diverse protein sources, including proteins associated with cancer metastasis and apoptosis regulation ^46,47,49^. These findings suggest that HLA-E may provide a more extensive monitor of cellular states than previously thought, which could enable surveillance by NK and potentially T cells of different cellular processes beyond MHC class I expression. Future studies will address under which physiological situations these peptides affect effector responses, whether they might constitute a substrate for altered-self recognition, and potential for involvement in tolerance or autoreactivity.

Characterization of the HLA-E and CD94/NKG2x repertoire enabled the identification of peptides capable of selectively activating NK cells through NKG2C, without inhibiting through NKG2A. Given the important role of CD94/NKG2x receptors in controlling NK cell function, and the association of adaptive NKG2C^+^ NK cells with viral control^9–11^ and anti-leukemic effects ^12^, these findings present new opportunities for therapeutic modulation of NK cell activity. Additionally, since HLA-E has been observed to be upregulated in cancer ^57^, these selectively activating peptides could be of particular interest for expanding NK cells in a therapeutic setting, without causing inhibition through CD94/NKG2A.

Finally, previous studies have identified T cell responses to MHC-E presenting peptides without obvious shared features ^19^. Through stability assays, we observed that activity-skewed peptides exhibit less HLA-E stabilizing effects, suggesting NKG2C has a higher tolerance for less stable pHLA-E, and the stability or conformation of HLA-E may play a role in activating T cells or NK cells.

## Methods

### Yeast-displayed pMHC design

Yeast-displayed HLA-E*01:03 was generated as a single chain trimer, covalently linked to a peptide and human β2M. Specifically, the C-terminus of the peptide was linked to N-terminus of β2M via Gly-Ser linker containing a Myc epitope tag (EQKLISEEDL) and protease site (LEVLFQGP). A second Gly-Ser linker connected the C-terminus of β2M to the N-terminus of HLA-E heavy chain α1, α2, and α3 domains. HLA-E contained a Y84A mutation to open the peptide binding groove to accommodate the peptide linker. This mutation could affect the binding preferences of the MHC F-pocket, but has been shown to largely recapitulate the endogenous binding properties of wild type class I MHCs ^27^. HLA-E C-terminus is connected to Aga2 via a final Gly-Ser linker containing an epitope tag (HA in clonal constructs: YPYDVPDYA; FLAG in library: DYKDDDDK). Yeast display constructs were generated in the pYAL vector, with the Aga2 leader sequence. Yeast strains were grown to confluence at 30°C in pH 5 SDCAA yeast media then subcultured into pH 5 SGCAA media at OD_600_ = 1.0 for 48-72 hours induction at 20°C ^58^.

### Tetramer and antibody staining yeast

To stain peptide-HLA-E (pHLA-E) with CD94/NKG2x tetramers, each biotinylated receptor was mixed with streptavidin coupled to AlexaFluor647 (made in-house) at a 5:1 ratio and incubated for 5 min on ice. Yeast were stained with 500 nM tetramer for 2 hours at 4°C in the dark with rotation. Then, the yeast were washed twice with FACS buffer (0.5% BSA and 2 mM EDTA in 1x PBS) before analysis via flow cytometry (Accuri C6 flow cytometer, BD Biosciences; Franklin Lakes, New Jersey).

Antibody staining was performed on yeast washed into FACS buffer, with antibody at a 1:50 or 1: 100 volume ratio. Yeast incubated with antibody for at least 20 minutes and excess antibody was removed by washing with FACS buffer. Staining was assessed on the Accuri C6 flow cytometer. Antibody clone 3D12 was used for anti-HLA-E staining.

### Yeast column enrichment competition assay

A yeast column enrichment competition assay was performed to evaluate pHLA-E binding to CD94/NKG2A receptor. Induced yeast expressing pHLA-E SCT were stained with an anti-epitope tag antibody (HA-488), then mixed with uninduced, unstained yeast at 1:50 ratio (2 x 10^5^ induced, 10^7^ uninduced). A sample of the mixture was assessed via flow cytometry analysis. The remaining mixed population of yeast were incubated with high avidity CD94/NKG2A-coated magnetic beads and enriched, as in library selections. The enriched population was assessed by flow cytometry. The HA-488 epitope stain differentiated pHLA-E-expressing yeast from uninduced competitor yeast. Representative gating is shown in **Supplemental Figure 12A**.

### Library design and selection

Randomized peptide libraries were generated using polymerase chain reaction (PCR) on the pMHC construct with primers encoding NNK degenerate codons (N = any base; K = G or T), to encode 9 or 10 randomized amino acids.

Randomized pMHC PCR product was mixed with linearized pYAL vector backbone at a 5:1 mass ratio and electroporated into electrocompetent RJY100 yeast ^59^. The final 9mer and 10mer libraries contained approximately 1.5 × 10^8^ and 7 × 10^7^ yeast transformants, respectively.

As described above, yeast were subject to selections for CD94/NKG2x binding. Selections were performed using streptavidin-coated magnetic beads (Miltenyi Biotec; Bergisch Gladbach, Germany) coupled to biotinylated CD94/NKG2x. Yeast were cultured, induced, and selected in three iterative rounds of selection. In a fourth round of selections, yeast were incubated with tetrameric AlexaFluor647-conjugated streptavidin (produced in-house) coupled to biotinylated CD94/NKG2x, and selected with ∝-AlexaFluor647 magnetic beads (Miltenyi Biotec). Each round was preceded by negative selection clearance round with uncoated streptavidin beads (rounds 1-3) or streptavidin and ∝-AlexaFluor647 beads (round 4). In round four, prior to addition of beads and selection, a sampling of CD94/NKG2x tetramer-stained yeast were assessed via flow cytometry on an Accuri C6 flow cytometer.

To prevent contamination with clonal yeast, the library template DNA encoded a stop codon in the peptide-encoding region. Additionally, the C-terminal HA tag was swapped for a FLAG tag to readily assess library expression and contamination.

### Library sequencing and analysis

Following selections, plasmid DNA from ~10^7^ yeast were extracted using Zymoprep II Yeast Miniprep Kit (Zymo Research; Irvine, CA) from each round of selection and the unselected library. Amplicons for deep sequencing were generated by PCR using the purified plasmids from yeast miniprep. Two rounds of PCR were completed to add homology for sequencing primers, i5 and i7 paired-end handles, and sequencing barcodes that are unique to each round of selection and selection reagent, to enable pooling of DNA. Amplicons were sequenced on an Illumina MiSeq (Illumina; San Diego, CA) with 2 x 150 nt paired-end reads at the MIT BioMicroCenter.

Paired-end reads were assembled and filtered for length and correct flanking sequences using PandaSeq ^60^. To correct for PCR or sequencing errors, and given the immense sampling space of peptides, sequences were clustered with more frequent sequences within Hamming Distance = 1 in DNA space using CDHit ^61,62^. Peptide sequences were translated from DNA and stop codon-containing sequences removed using an in-house script.

**Supplemental Data 1** contains processed peptide data. In this file, sequences not containing stop codons are listed with their read counts, labeled for round of selection and selection reagent. In the column header, the letter indicates the selection reagent, and number indicates the selection round (e.g. “A1” is post-round 1 of selection with NKG2A). “R0” is the unselected round 0 library, and cross-selected libraries are labelled with their original selection reagent followed by the current selection reagent (e.g. “CA4” is post-round 4 of selection using NKG2A on the library that was previously selected with NKG2C).

### Heat map and sequence logo visualization of library-enriched peptides

Sequence logo visualizations were generated using the sequence logo generator WebLogo ^63^. Sequences were repeated *int(count/100)* times, which filters possible noise with read counts <100 and weights sequences by frequency. Sequence logos were generated using WebLogo default settings.

Heatmaps were generated with a custom script, on all enriched peptides from a given round of selection. Frequencies were calculated by weighting each peptide by its read count. Maps were binned as shown to capture the frequency of reads that encode a peptide with a given amino acid at a given position.

### Clustering peptides to identify subdominant motifs

We identified the subdominant cluster of WN peptides using various clustering methods, including Gibbs Cluster 2.0 ^64^ on peptides enriched in Round 3 of CD94/NKG2A selections (not weighted by read counts), excluding stop codon-containing peptides, using the default method “MHC class I ligands of same length”, modified to assess 1-15 clusters, with the trash cluster to remove outliers turned off.

### Prediction algorithm generation

CD94/NKG2A- or CD94/NKG2C-specific prediction models were trained on yeast display library data, with positive examples drawn from Round 3 of selections, with counts greater than or equal to 100, each appearing in the training data *int(count/100)* times. Negative examples were selected from the unselected library, with read count equal to 1 and excluding stop codon-containing peptides. The total number of negative peptides were selected such that there were equal numbers of positive and negative examples in the training set (total number, not unique examples). Positive examples were assigned a target value of 1 and negative examples were assigned a target value of 0.

These data were used to train NNAlign 2.0 with MHC Class I defaults, excepting Maximum Length for Deletions, Maximum Length for Insertions, and Length of PFR for Composition Encoding were set to zero; Encode PFR Length was set to −1 (for no encoding); and both Binned Peptide Length Encoding and Length of Peptides Generated from FASTA Entries were set to 9.

### Prediction on proteome-derived peptides

With a 9mer window, step size 1, 11,019,710 unique human proteome 9mers were generated from a reference proteome (Uniprot UP000005640), excluding peptides containing selenocysteine. The same was done for human CMV (reference Uniprot proteome UP000000938), generating 61,303 peptides. UL120_71-79, AD169_ and UL120_71-79, BE/33/2010_ were derived from sequences in Uniprot UP000008991 and UP000100992, respectively.

A one-sided Mann Whitney U test was performed on HLA-E-binding VL9 and VL9-like signal peptides from the human proteome compared to other 9mer human proteome peptides (11,019,710 total 9mer peptides). *U*-values were 66,115,459 (NKG2A) and 66,118,060 (NKG2C) and were calculated alongside *p*-values using scipy version 1.4.1.

### NetMHC predictions

NetMHCpan4.0 predictions were performed with a local version of the algorithm on 9mer peptides for HLA-E*01:03.

### Recombinant protein production

Recombinant soluble HLA-E single chain trimers, CD94/NKG2A, and CD94/NKG2C for SPR were produced using a baculovirus expression system with High Five (Hi5) insect cells (ThermoFisher). Individual constructs were cloned into pAcGP67a vectors. For each, 2 μg of plasmid DNA was transfected into SF9 insect cells using BestBac 2.0 baculovirus DNA (Expression Systems; Davis, CA) and Cellfectin II reagent (ThermoFisher). Viruses were propagated to high titer and transduced into Hi5 cells, grown at 27°C for 48–72 hours, and purified from pre-conditioned cell media supernatant using Ni-NTA resin and size exclusion chromatography with a S200 column on an AKTAPure FPLC (GE Healthcare; Chicago, IL).

Single chain timers were formatted with the C-terminus of the peptide linked to the N-terminus of human β2M via a flexible linker. In turn a flexible Gly-Ser linker connects the C-terminus of β2M to the N-terminus of the extracellular α1, α2, and α3 domains of HLA-E*01:03 heavy chain, containing a Y84A mutation to accommodate the linked peptide. An AviTag biotinylation tag and poly-histidine tag were connected to the C-terminus of HLA-E heavy chain. Protein was biotinylated overnight before FPLC-based purification.

CD94/NKG2x proteins were expressed as single-chain fusion proteins. The C-terminus of human NKG2A or NKG2C ectodomain was connected via a GGSGGS linker to human CD94 ectodomain. The C-terminus of CD94 is connected to an AviTag biotinylation tag and poly-histidine tag. Protein for SPR was used fresh and dialyzed overnight into HBS-EP+ buffer (10 mM HEPES pH 7.4, 150 mM NaCl, 3 mM EDTA and 0.05% v/v Surfactant P20) (Cytiva; Malborough, MA) or buffer exchanged into HBS-EP+ buffer during FPLC-based purification before use in SPR. CD94/NKG2x for selections was expressed from the Expi293 expression system (ThermoFisher; Waltham, MA) as per the manufacturer’s recommendations, using a modified pHLSec-Avitag3-His6 vector ^65^, and purified by IMAC (HisTrap Excel column) and SEC (S200 Increase GL 10/30) utilizing an Akta Pure (Cytiva). Biotinylation was confirmed by streptavidin gel shift assay ^66^.

### Surface plasmon resonance

Steady-state surface plasmon resonance experiments were performed with a Biacore T200 instrument. CD94/NKG2x in HBS-EP+ buffer was injected as analyte in a concentration range of 0.1 μM to 102.4 μM and flow rate of 10 μL/min at 25°C. Data was fit with Prism 9.4 (GraphPad Software Inc; San Diego CA) to a “one site, specific binding” model. Sensorgrams are included in **Supplemental Figure 11**.

For investigating WN peptides, biotinylated pMHC was immobilized at approximately 400 Response Units (RU) on a Series S SA sensor chip (Cytiva). WN peptide HLA-E SCT, VL9 peptide HLA-E SCT, and reference HLA-DR401 (HLA-DRA1*01:01, HLA-DRB1*04:01, linked to CLIP peptide) ^31^ were immobilized. CD94/NKG2A was injected followed by CD94/NKG2C. Approximate K_D_ values calculated by Biacore software are as follows: WN SCT + CD94/NKG2A: 41.5 μM; WN SCT + CD94/NKG2C: 17.4 μM; VL9 SCT + CD94/NKG2A: 8.2 μM; VL9 SCT + CD94/NKG2C: 21.4 μM.

For investigating human-derived peptides, Series S CM5 chips (Cytiva) were coupled to neutravidin and utilized. Biotinylated pMHC was immobilized at approximately 400 RU. On one chip, VL9 HLA-E SCT, BFAR_263-271_ HLA-E SCT, CREB3L1_419-427_ HLA-E SCT, and reference HLA-DR401 were immobilized. CD94/NKG2A was injected followed by CD94/NKG2C. On a second chip, PISD55-63 HLA-E SCT, AKAP6_388-396_ HLA-E SCT, GTF3C5_293-301_ HLA-E SCT, and reference HLA-DR401 were immobilized. CD94/NKG2C was injected followed by CD94/NKG2A. K_D_ values calculated by Biacore software are reported in **Figure 5**. R_max, fit_ values were calculated by Biacore software. R_max, theoretical_ values were calculated using the analyte/ligand mass ratio (0.66) multiplied by the amount of ligand coupled to the chip.

### Prometheus differential scanning fluorimetry experiments

Differential scanning fluorimetry (DSF) experiments were performed with a Prometheus NanoTemper NT.48 (Munich, Germany) to measure protein stability, using intrinsic tryptophan fluorescence.

For these experiments, refolded empty HLA-E was generated. Human β2M plus a polyhistidine tag and HLA-E*01:03 extracellular α1, α2, and α3 domains plus an AviTag biotinylation tag were separately codon optimized for *E. coli* and cloned into the bacterial expression vector pET28a. Inclusion bodies for HLA-E and β2M were made in BL21 *E. coli*. Inclusion bodies were purified, homogenized, and dissolved, including denaturation in 8 M urea solution. Before refolding, each IB was mixed with an equal volume of Gdn-Cl solution (6 M Guanidine HCl, 500 mM Tris, 2 mM EDTA, 100 mM NaCl) and incubated overnight at 37°C. β2M was injected dropwise with a 27-gauge needle into refolding buffer (100 mM pH 8.3 Tris, 400 mM L-arginine hydrochloride, 2 mM EDTA, 0.5 mM oxidized glutathione, 5 mM reduced glutathione, 0.2 mM PMSF) and incubated, stirring gently, at 4°C for 1 hour, after which an equal mass of HLA-E heavy chain was similarly injected. This solution incubated at 4°C overnight, followed by two days of dialysis in 10 mM Tris + 50 mM NaCl. Protein was concentrated in 10 kDa molecular weight cutoff centrifugal concentrators and purified by size exclusion chromatography on an AKTAPURE FPLC (GE Healthcare, Chicago IL), using an S200 column followed by a S75 column. Protein was concentrated with size filters as initial attempts to concentrate using the poly-histidine tag and Ni-NTA resin resulted in the protein precipitating out of solution.

Individual DSF reactions were set up with 9 μg of MHC and 100x, 50x, or 10x peptide to MHC ratio by molarity (peptides from GenScript). Final reaction volume was 20 μL and reaction mixtures were incubated at room temperature for 20-30 minutes.

Prometheus excitation power was set such that raw fluorescence counts were between 8000-15000 for each sample. Samples were heated at 1°C per minute, from 20-95°C. DSF measurements were taken in duplicate with high-sensitivity and standard capillaries. Melting temperatures were similar for a given peptide across conditions, and representative data from 50x ratio in high sensitivity capillary are presented. A secondary inflection point around 60°C in most melts is consistent with melting of β2M ^67^.

### Cell lines

K562/HLA-E ^45^ and RMA-S/HLA-E ^52^ were maintained in complete medium (RPMI-1640 containing glutamine and supplemented with 10% v/v FBS, 20□μM β-mercaptoethanol and 100 U/mL penicillin-streptomycin; all Thermo Fisher) in the presence of 400□μg/mL hygromycin B and 1□mg/mL G418 (both InvivoGen), respectively.

### HLA-E surface stabilization assay

RMA-S/HLA-E cells were incubated with serial dilutions of peptides (3-300 μM, Genscript) in OptiMEM (ThermoFisher) for 16 h at 37°C. Cells were washed with PBS, stained for HLA-E (Clone 3D12, Biolegend) for 15 min at room temperature and analyzed by flow cytometry. Peptide UBAC2_275-279_ was ordered as a 5mer peptide (WNRLF) given the similarity with other WN peptides and hypothesis of a truncated peptide binding in the HLA-E groove. Due to lower peptide solubility, the maximal concentration tested for BFAR_263-271_ was 150 μM, for EMC1_725-733_ was 75 μM, for OR5D14_88-96_ was 30 μM, and for UL102_292-300_ was 30 μM. Measurements from replicate experiments are plotted in **Figure 3** and **Supplemental Figure 7.**

### NK cell activation assays

All analyses of human data were carried out in compliance with the relevant ethical regulations. Healthy blood donors gave informed consent at DRK Blutspendedienst Nord-Ost, Dresden, Germany, and buffy coats were obtained as approved by Charité ethics committee (EA4/059/17). PBMCs were isolated from human CMV-seropositive buffy coats by density centrifugation over Ficoll Paque Plus (GE Healthcare) and screened for the presence of NKG2C^+^ NK cell expansions by flow cytometry ^51^. NK cells from donors with NKG2C^+^ expansions were enriched with CD56 microbeads (Miltenyi Biotec) and cryopreserved in FBS with 10 % v/v DMSO. After thawing, NK cells were washed with complete medium, stained with antibodies and Fixable Viability Dye eFluor 780 (Thermo Fisher) for 15 min at 4°C in PBS, sorted as viable CD56^+^ CD3^-^ cells and rested in complete media overnight. RMA-S/HLA-E or K562/HLA-E were pulsed with the indicated peptides at a concentration of 300 μM as described above (150 μM for BFAR_263-271_ due to lower peptide solubility). NK cells were co-cultured with target cells at an effector-to-target ratio of 2:1 in complete media in the presence of 300 μM peptide (150 μM for BFAR_263-271_) and anti-CD107a. Brefeldin A and Monensin (BD Biosciences) were added after 1 h culture period. After an additional 5 h, the stimulation was stopped by centrifugation at 4°C. Cells underwent surface staining, followed by fixation and intracellular staining using Inside Stain Kit (Miltenyi Biotec). Samples were analyzed by flow cytometry on an LSR Fortessa (BD Biosciences). **Supplemental Figure 12B** shows the gating strategy used for NK cells. Antibodies are listed in **Supplemental Table 1**.

## Data Availability

All deep sequencing data are available on the Sequence Read Archive (SRA) with accession code PRJNA859187 [http://www.ncbi.nlm.nih.gov/bioproject/859187]. Processed peptide data are provided in **Supplemental Data 1**.

## Code Availability

Scripts for data processing and analysis and NNAlign model files are publicly available at https://github.com/birnbaumlab/Huisman-et-al-2022-HLA-E.

## Acknowledgements

We would like to thank Lucy Walters for helpful discussions about HLA-E, Max Quastel for generously sharing HLA-E Prometheus protocols, and the MIT BioMicro Center for library sequencing. This work was supported in part by the Koch Institute Support (core) Grant P30-CA14051 from the National Cancer Institute. This work was supported by National Institute of Health (U19-AI110495), the Melanoma Research Alliance Foundation, and the Packard Foundation to M.E.B; a National Science Foundation Graduate Research Fellowship to B.D.H; Deutsche Forschungsgemeinschaft (DFG) grants SPP 1937 (RO3565/4-2) and SFB TRR241 B02 to C.R.; Leibniz-Science Campus Chronic Inflammation and Leibniz-Kooperative Exzellenz K259/2019 to C.R.; and Berlin Health Innovations (BHI) Validation Fund to C.R. and T.R..

## Author Contributions

Project conception: B.D.H., N.G., M.E.B.; Conducted experiments: B.D.H., N.G., T.R., L.G., N.K.S.; Performed data analyses: B.D.H., N.G., T.R.; Supervised the work: A.J.Mc.M., G.G., C.R., M.E.B.; Wrote the manuscript: B.D.H., M.E.B.; All authors contributed to the editing of the manuscript.

