## Supplemental Figures for "An unbiased characterization of the HLA-E and CD94/NKG2x peptide repertoire reveals peptide ligands that skew NK cell activation"

**12 Supplemental Figures**
**1 Supplemental Table**

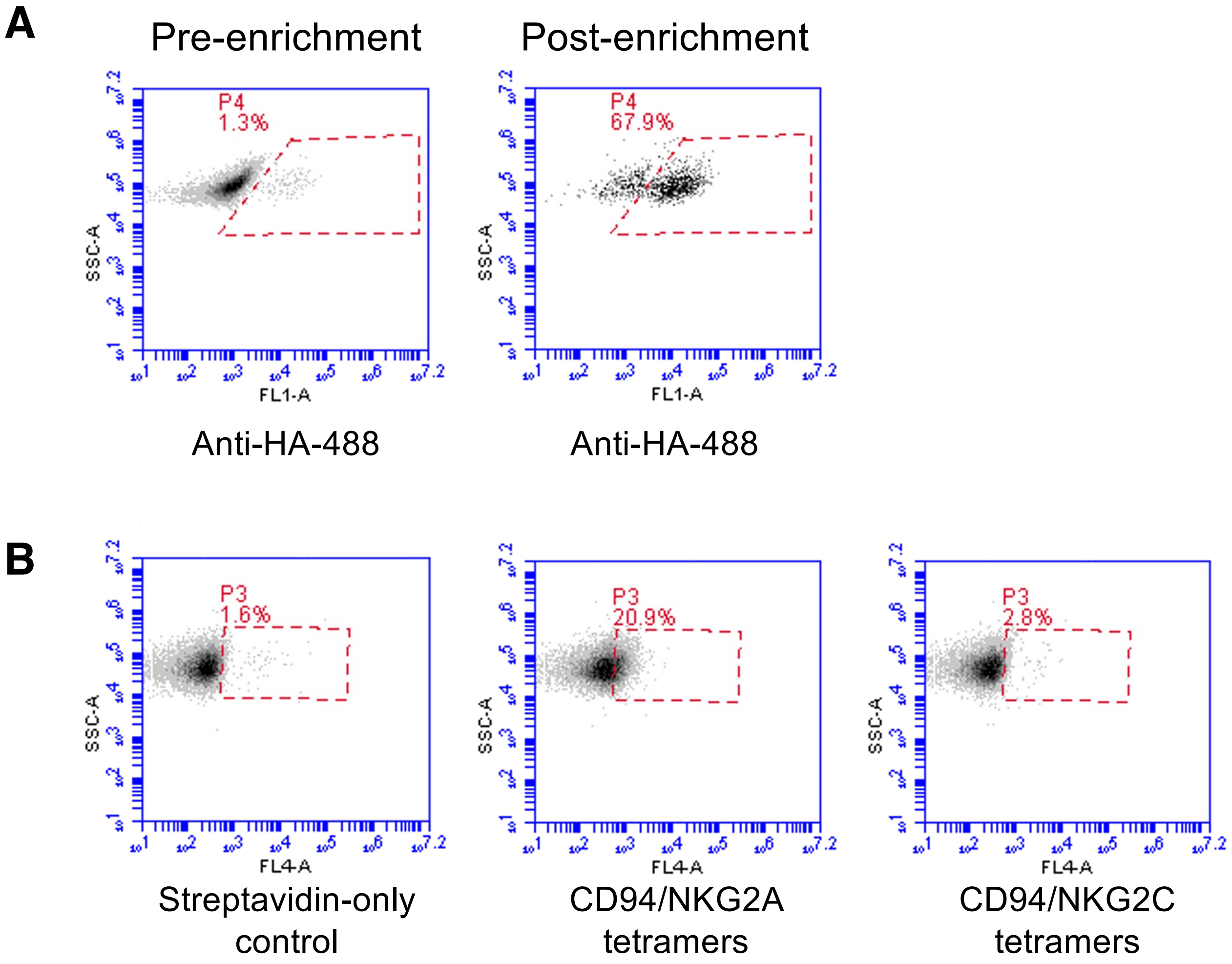


**Supplemental Figure 1. Validation of HLA-E construct. a)** Column enrichment assay of VL9-HLA-E with CD94/NKG2A. Stained yeast are doped into an uninduced yeast population at 1:50 mixing ratio and enriched with CD94/NKG2A-coated beads, and the fraction of stained yeast are shown pre- and post-enrichment. **b)** Validation staining of HLA-E with VL9 peptide with CD94/NKG2A (same as **Figure 1**) and CD94/NKG2C tetramers, made with Streptavidin-647, with Streptavidin-647-only control.

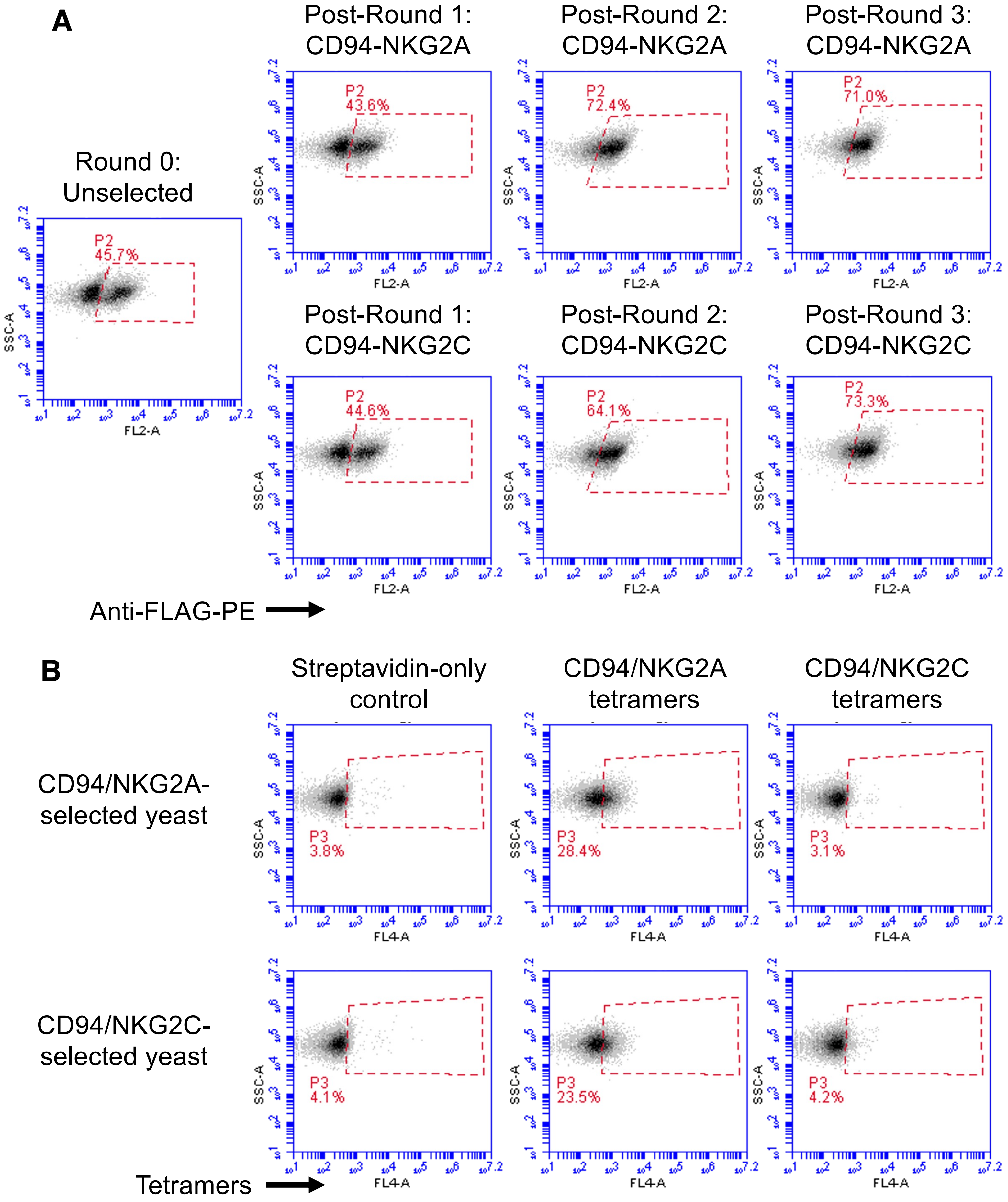


**Supplemental Figure 2. Library dynamics over rounds of selection with CD94/NKG2A or CD94/NKG2C. a)** Prior to selections, a sampling of yeast were assessed for FLAG epitope tag expression, which increased over rounds of selection (gate drawn daily on unstained yeast). **b)** Staining of yeast during Round 4 with CD94/NKG2A or CD94/NKG2C tetramers made with Streptavidin-647, on libraries previously selected with CD94/NKG2A or CD94/NKG2C in Rounds 1-3, including negative Streptavidin-647-only control.

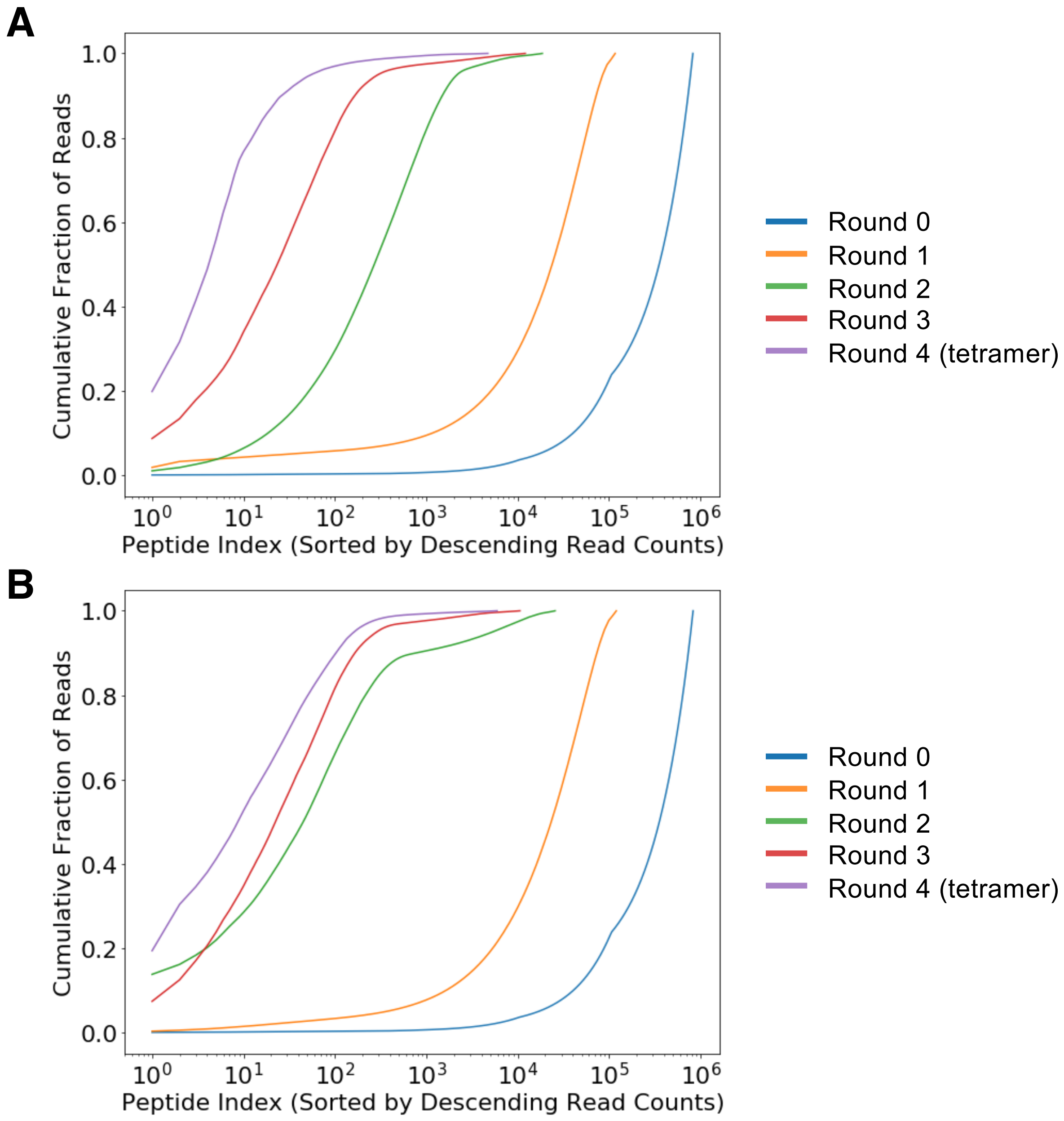


**Supplemental Figure 3. Peptide frequency in deep sequencing data.** Cumulative fraction of read counts for peptides selected with **a)** CD94-NKG2A or **b)** CD94-NKG2C.

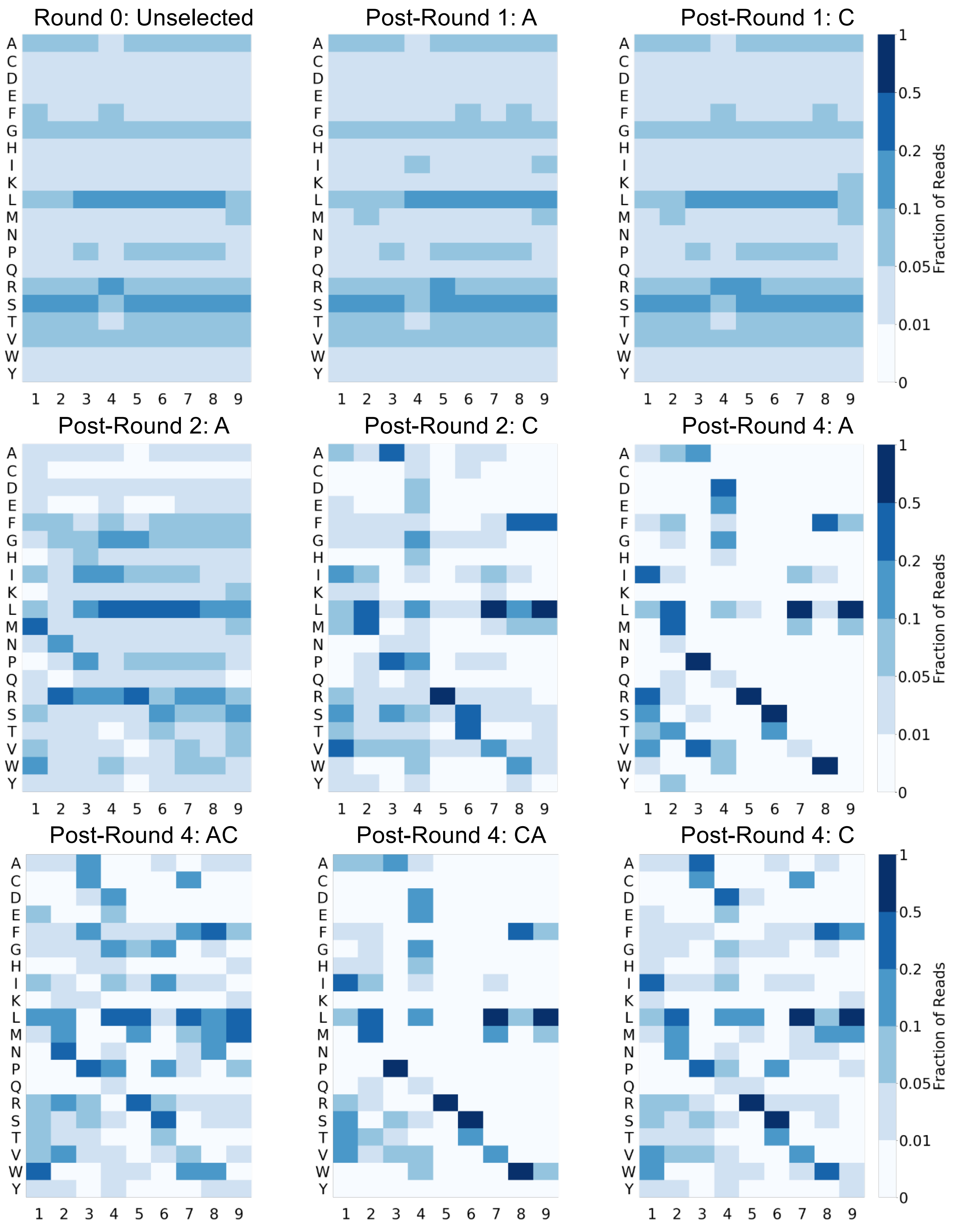


**Supplemental Figure 4. Heatmaps for other rounds of selection.** A: Selected with CD94/NKG2A; C: Selected with CD94/NKG2C; AC: yeast previously selected with CD94/NKG2A, cross-selected with CD94/NKG2C in this round; CA: yeast previously selected with CD94/NKG2C, cross-selected with CD94/NKG2A in this round.

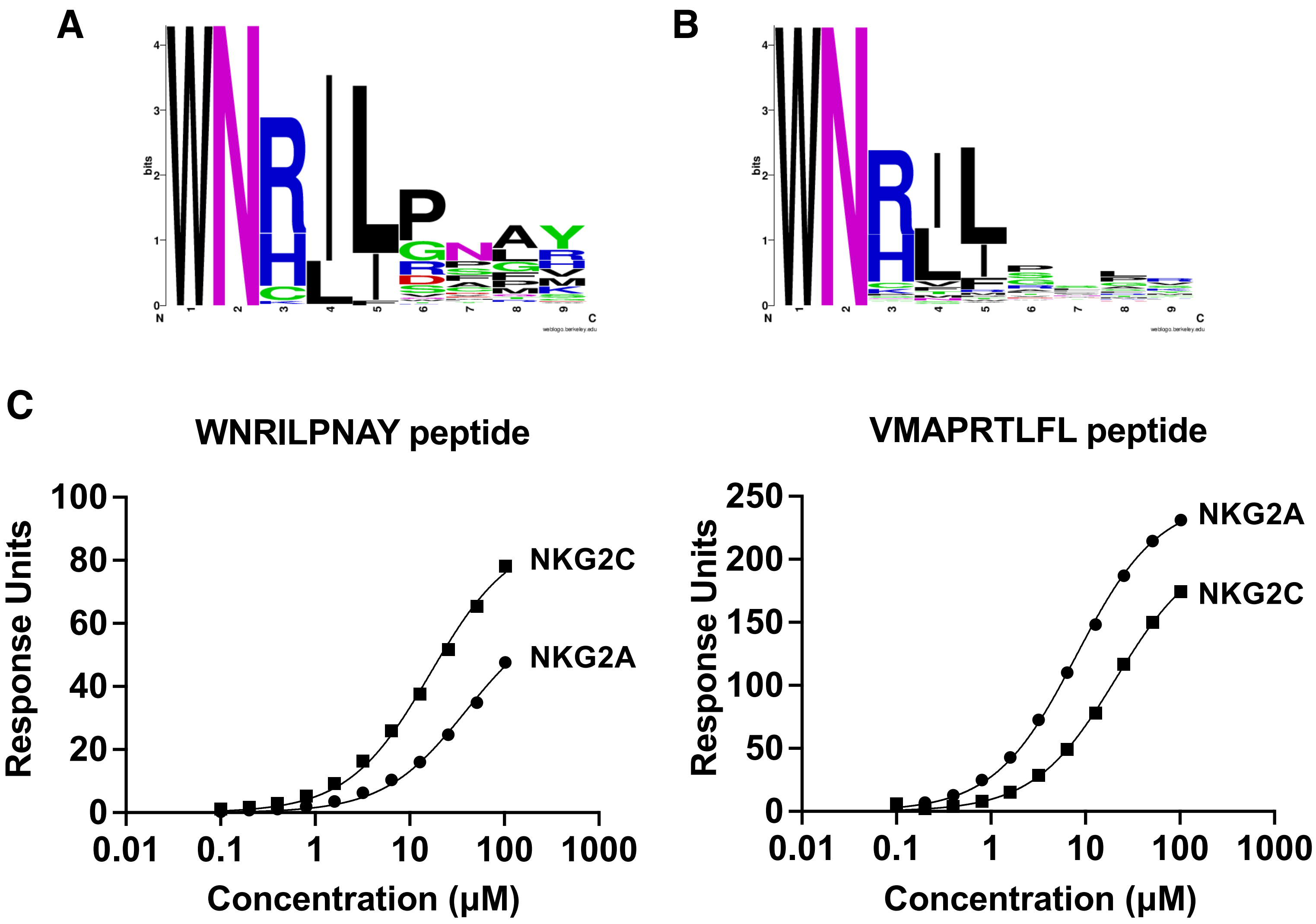


**Supplemental Figure 5. Subdominant motif of peptides with P1 Trp and P2 Asn (WN peptides).** Sequence motifs of peptides that have P1 Trp and P2 Asn, **a)** weighted by frequency or **b)** not weighted by frequency in Round 3 of selections with CD94/NKG2A. **c)** Surface plasmon resonance data for HLA-E single chain trimers containing WN 9mer peptide (left) or VL9 control peptide (right), binding to CD94/NKG2A or CD94/NKG2C.

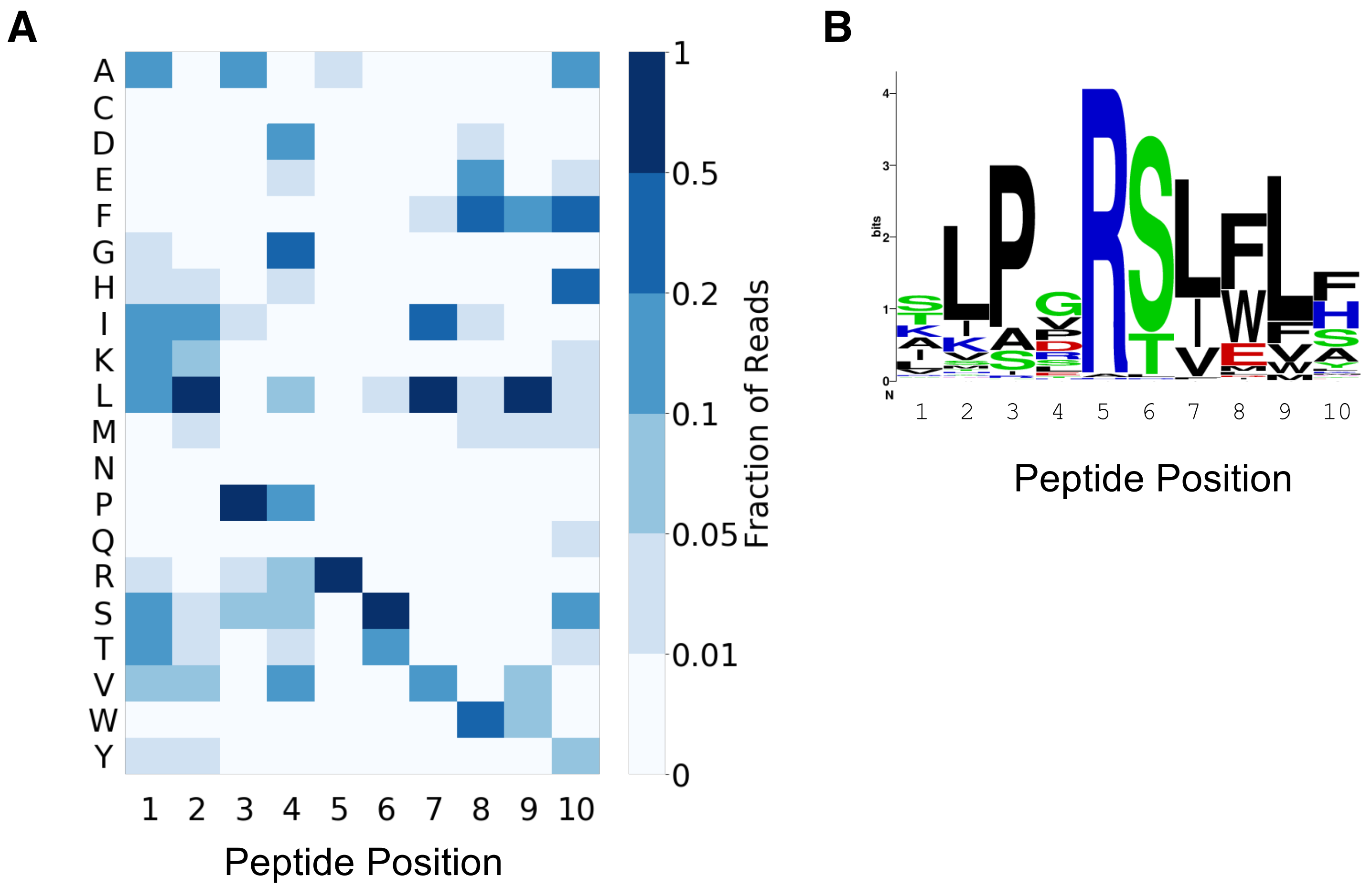


**Supplemental Figure 6. 10mer peptide data a)** Heatmap and **b)** sequence logo representation of peptide positional amino acid preferences in 10mer peptides after three rounds of selection with CD94/NKG2C, with peptides weighted by read count.

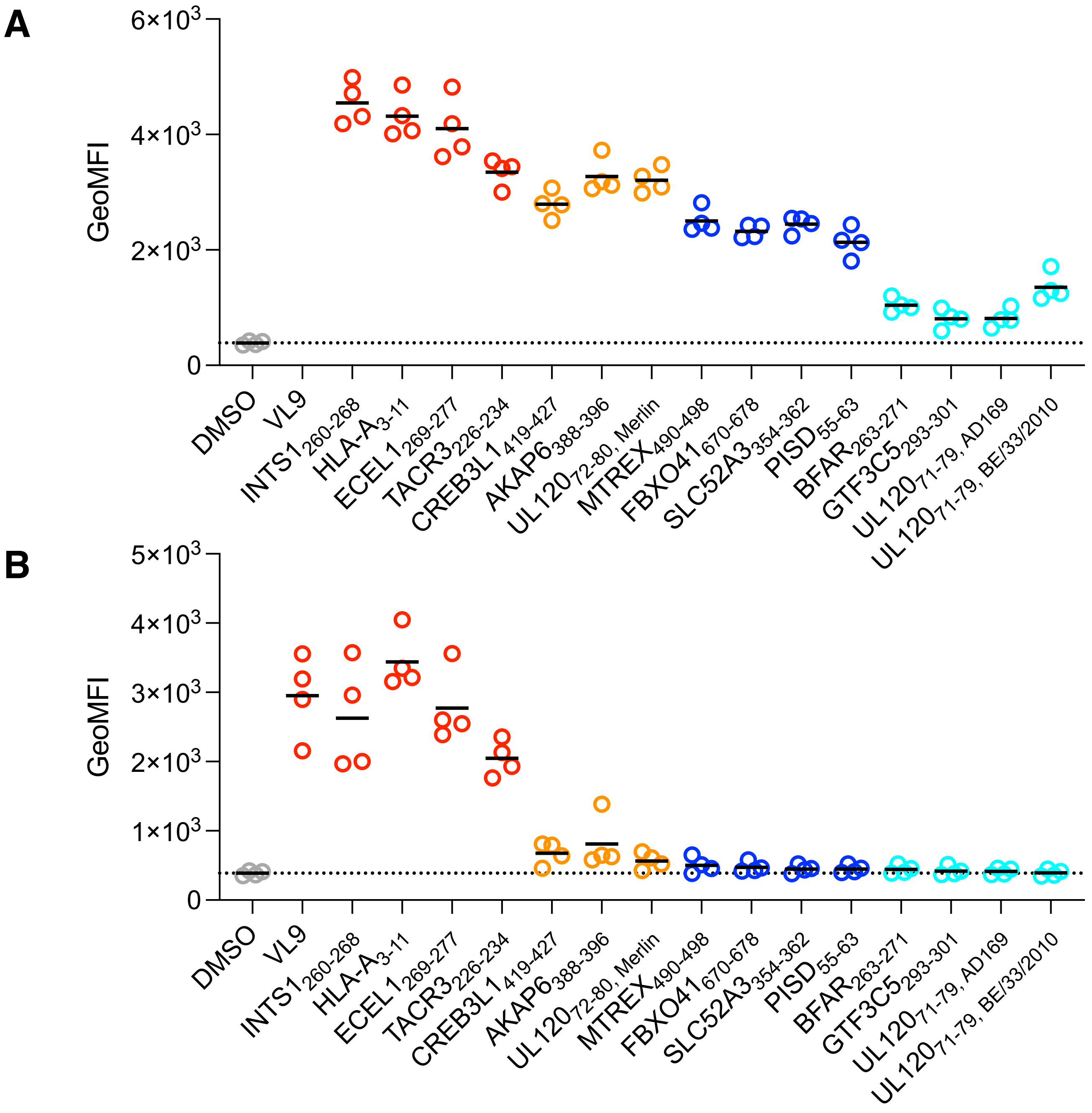


**Supplemental Figure 7. Peptide stabilization of HLA-E surface expression at additional concentrations**. Assessment of peptide-HLA-E binding via HLA-E surface stabilization assay with RMA-S/HLA-E cells incubated with **A)** 300 µM peptide (150 µM for $BFAR_{263\text{-}271}$ due to lower peptide solubility) and **B)** 3 µM peptide. Measurements from replicate experiments are shown, with solid black lines indicating mean values.

A

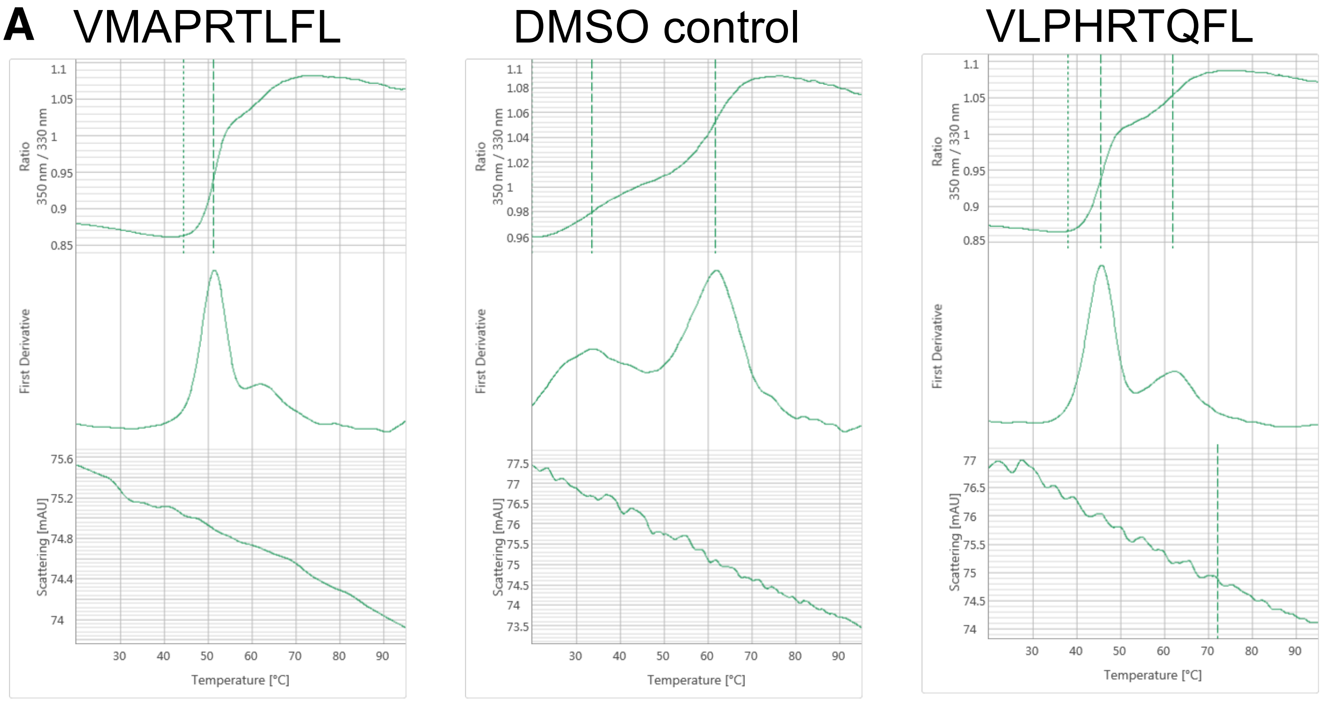


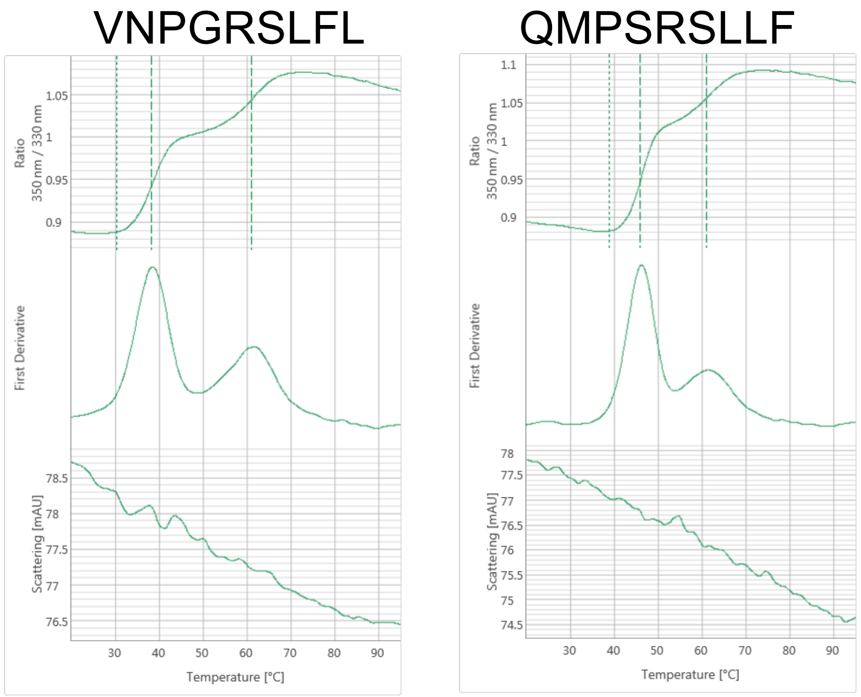


B

| Peptide (Name) | $T_m$ (°C) |
|---|---|
| VMAPRTLFL (LFL) | 51.5 |
| QMPSRSLLF (CREB3L1$_{419\text{-}427}$) | 46.2 |
| VLPHRTQFL (UL120$_{72\text{-}80,\ Merlin}$) | 45.7 |
| VNPGRSLFL (BFAR$_{263\text{-}271}$) | 38.5 |
| DMSO control | 33.8 |

**Supplemental Figure 8. Differential scanning fluorimetry data generated via Prometheus NanoTemper for predicted peptides with HLA-E. a)** Raw 350/330 nm ratios and first derivative for samples. The second inflection point is consistent with β2M unfolding. **b)** Summary of $T_m$ values for the first inflection point called by Prometheus software for these samples.

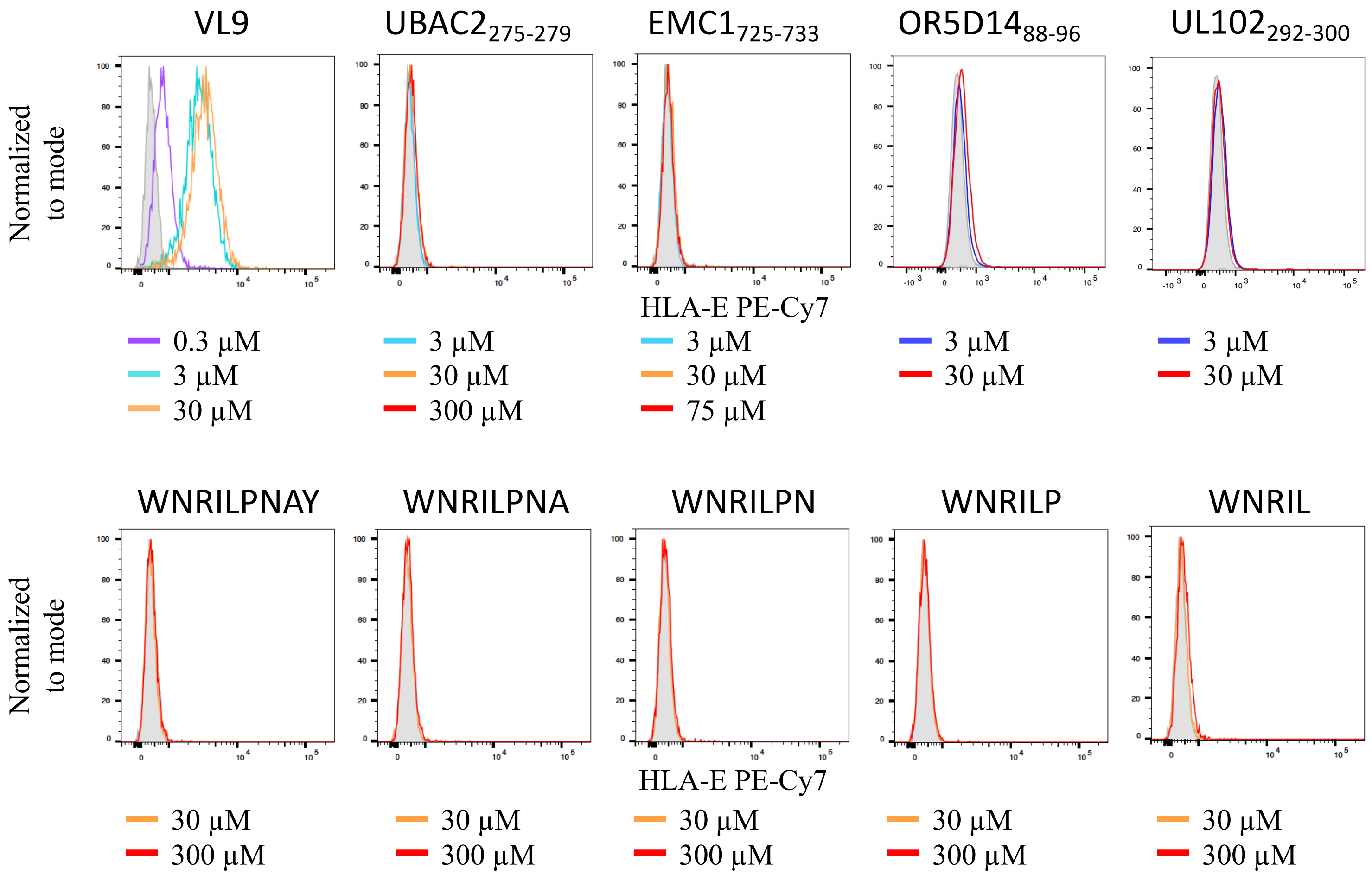


**Supplemental Figure 9. Stabilization data for peptides which did not stabilize HLA-E surface expression.** Peptides UBAC2$_{275\text{-}279}$, EMC1$_{725\text{-}733}$, and OR5D14$_{88\text{-}96}$, and UL102$_{292\text{-}300}$ were assessed, along with length variants of WN peptides. VL9 was included as a control. Peptide concentrations tested are indicated, where maximum concentrations assessed vary due to differing peptide solubilities. Grey shaded curve is DMSO control.

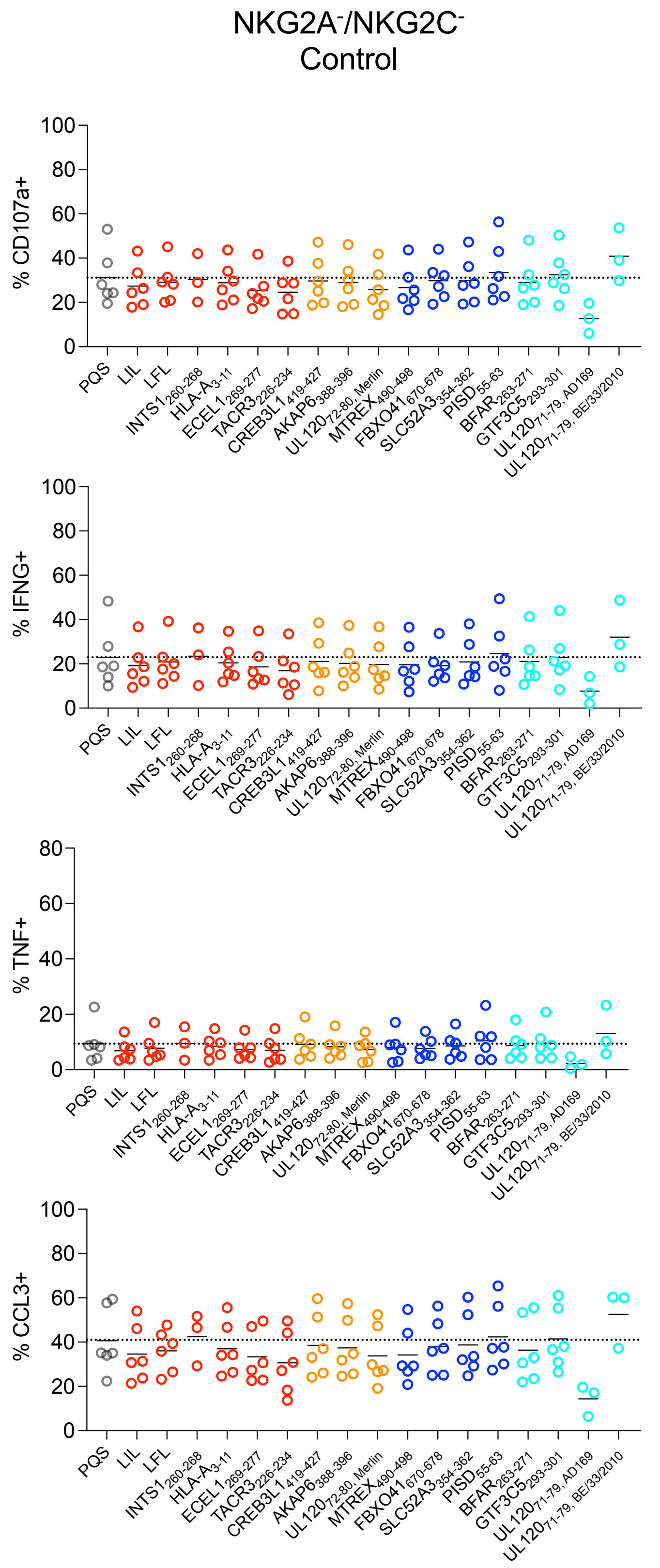


**Supplemental Figure 10. NKG2A$^-$/NKG2C$^-$ NK cell controls.** Assessment of affects of peptides on control NKG2A$^-$/NKG2C$^-$ NK cell activity after co-incubation with peptides and K562/HLA-E cells. Peptides were derived from the human or CMV proteomes or are included as positive (LFL, LIL) or negative controls (PQS). Replicates for individual peptides are from different donors, with solid black lines indicating mean values.

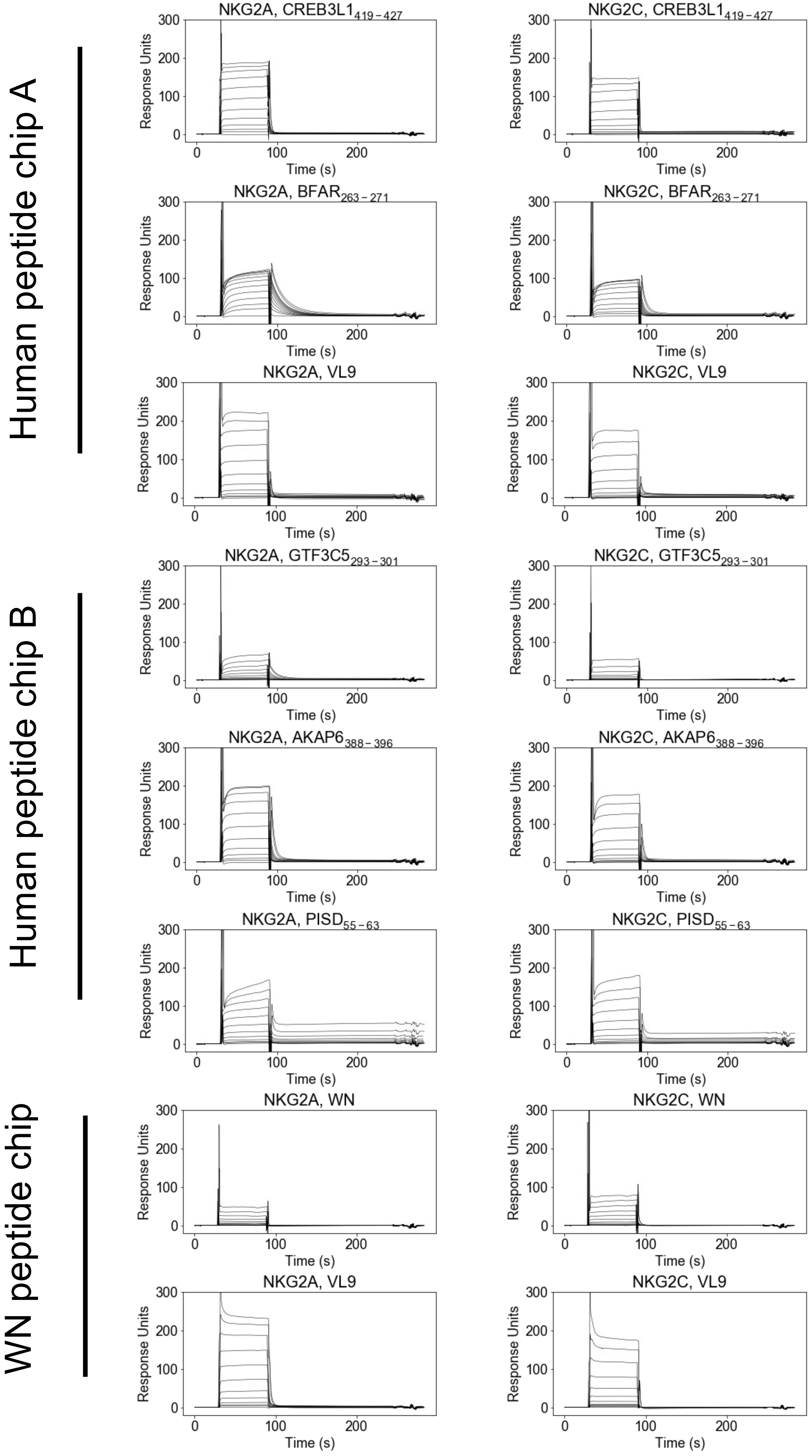


**Supplemental Figure 11. SPR sensorgrams.** Sensorgrams with peptide, receptor, and chip indicated.

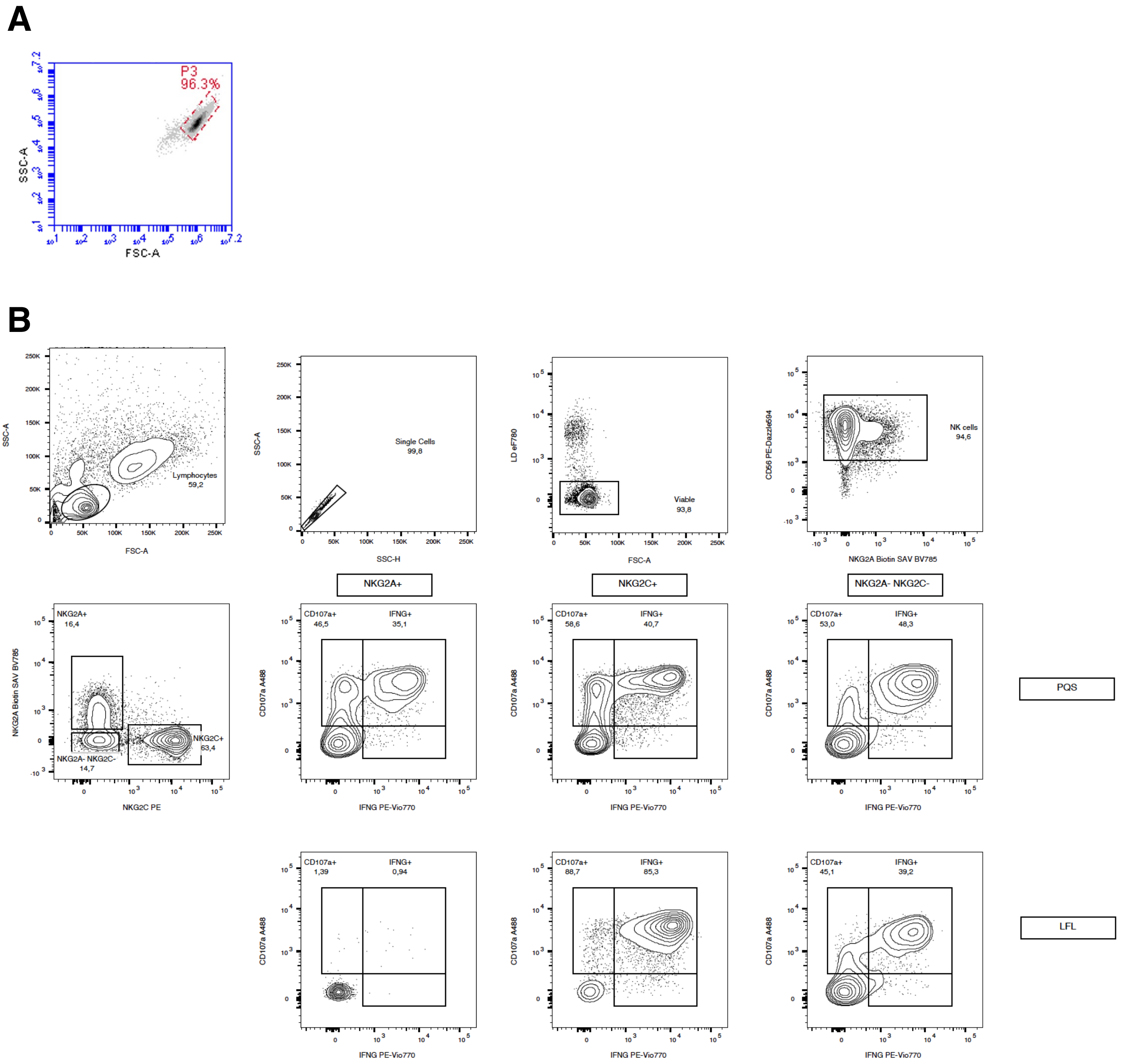


**Supplemental Figure 12. Flow cytometry gating. a)** Representative gate for yeast display column enrichment assay. **b)** Gating strategy for NK cell stimulation assays.

| Antibody | Source | Identifier |
|---|---|---|
| PE/Dazzle™ 594 anti-human CD56 | Biolegend | AB_2563564 (BioLegend Cat# 318348) |
| CD3 Monoclonal Antibody (SK7), APC-eFluor 780 | Invitrogen | AB_10717514 (ThermoFisher Cat# 47-0036-42) |
| CD159a (NKG2A) Antibody, anti-human, Biotin, REAfinity | Miltenyi Biotec | AB_2783969 (Miltenyi Biotec Cat# 130-114-090) |
| CD159c (NKG2C) Antibody, anti-human, PE, REAfinity™ | Miltenyi Biotec | AB_2751866 (Miltenyi Biotec Cat# 130-119-814) |
| Alexa Fluor® 488 anti-human CD107a (LAMP-1) Antibody | Biolegend | AB_1227504 (BioLegend Cat# 328610) |
| IFN-γ Antibody, anti-human, PE-Vio® 770, REAfinity™ | Miltenyi Biotec | AB_2652240 (Miltenyi Biotec Cat# 130-109-235) |
| Brilliant Violet 605™ anti-human TNF-α Antibody | Biolegend | AB_2563884 (BioLegend Cat# 502936) |
| CCL3 (MIP-1α) Antibody, anti-human, REAfinity™ | Miltenyi Biotec | AB_2651376 (Miltenyi Biotec Cat# 130-103-630) |
| PE/Cyanine7 anti-human HLA-E Antibody | Biolegend | AB_2565263 (BioLegend Cat# 342608) |
| PE anti-human HLA-E Antibody | Biolegend | AB_1659250 (BioLegend Cat# 342603) |
| Alexa Fluor ® 647 HA-Tag Mouse mAb | Cell Signaling Technology | Cell Signaling Technology Cat# 3444S |
| Alexa Fluor ® 488 HA-Tag Mouse mAb | Cell Signaling Technology | Cell Signaling Technology Cat# 2350S |
| PE DYKDDDDK Tag (D6W5B) Rabbit mAb | Cell Signaling Technology | Cell Signaling Technology Cat# 98533S |

**Supplemental Table 1.** Antibodies used in this work.
